# Dynamic Overrepresentation of Accumbal Cues in Food- and Opioid-Seeking Rats after Prenatal THC Exposure

**DOI:** 10.1101/2024.05.06.592839

**Authors:** Miguel Á. Luján, Reana Young-Morrison, Sonia Aroni, István Katona, Miriam Melis, Joseph F. Cheer

**Affiliations:** Department of Neurobiology, University of Maryland School of Medicine, Baltimore, MD, USA; Department of Biomedical Sciences, University of Cagliari, Cittadella Universitaria, Monserrato, Italy; Momentum Laboratory of Molecular Neurobiology, Institute of Experimental Medicine, Hungarian Academy of Sciences, Budapest, Hungary; Department of Psychological and Brain Sciences, Indiana University, Bloomington, IN, USA; Department of Psychiatry, University of Maryland School of Medicine, Baltimore, MD, USA

## Abstract

The increasing prevalence of cannabis use during pregnancy has raised significant medical concerns, primarily related to the presence of Δ9-tetrahydrocannabinol (THC), which readily crosses the placenta and impacts fetal brain development. Previous research has identified midbrain dopaminergic neuronal alterations related to maternal THC consumption. However, the enduring consequences that prenatal cannabis exposure (PCE) has on striatum-based processing during voluntary reward pursuit have not been specifically determined. Here, we characterize PCE rats during food (palatable pellets) or opioid (remifentanyl)-maintained reward seeking. We find that the supra motivational phenotype of PCE rats is independent of value-based processing and is instead related to augmented reinforcing efficiency of opioid rewards. Our findings reveal that in utero THC exposure leads to increased cue-evoked dopamine release responses and an overrepresentation of cue-aligned, effort-driven striatal patterns of encoding. Recapitulating findings in humans, drug-related neurobiological adaptations of PCE were more pronounced in males, who similarly showed increased vulnerability for relapse. Collectively, these findings indicate that prenatal THC exposure in male rats engenders a pronounced neurodevelopmental susceptibility to addiction-like disorders later in life.

## Introduction

Use of cannabis during pregnancy is on the rise^1,2^ with an estimated prevalence ranging from 7%, according to self-reports^3^, to 22.4% when analyzing umbilical cord samples^4^. Pregnant individuals report using cannabis to relieve stress and anxiety^5^, a trend worsened by the COVID-19 pandemic^6^. This alarming trajectory is compounded by misleading marketing strategies by medically-licensed dispensaries^7^, and a growing misconception about the safety of “natural” cannabis plant derivates^8^. The use of cannabis during pregnancy comes with significant medical concerns, particularly due to the presence of Δ^9^-tetrahydrocannabinol (THC), the main psychoactive component of cannabis. THC readily crosses the placenta, and a third of plasma content enters the fetus^9^. Exogenous cannabinoid receptor-type 1 (CB1R) agonists, like THC, interfere with endocannabinoid signaling cascades governing progenitor cell proliferation, neuronal differentiation, axon growth, synapse formation and pruning in the developing brain^10^.

Human longitudinal cohorts have underscored the detrimental effects of prenatal cannabis exposure (PCE), predisposing offspring to a range of neuropsychiatric conditions such as hyperactivity, impulsivity, and increased susceptibility to psychosis^11^. Rodent studies from our laboratory and others have shed light into the neurobiological disturbances caused by *in utero* THC exposure, consistently impacting brain dopaminergic pathways along the mesolimbic system^12^. In the ventral tegmental area (VTA), PCE leads to a maladaptive phenotype characterized by the hyperexcitability of dopamine neurons^13–15^. In the nucleus accumbens (NAc), where VTA dopamine neurons project^16^, maternal THC exposure induces enduring epigenetic and motivational signatures^17^. In light of these concerning findings, and human studies disclosing an increased risk of substance use ^18,19^, we hypothesize that exposure to THC during pregnancy enhances the reinforcing and dopaminergic effects of drugs of abuse later in life.

To test this possibility, this study examined reward processing as well as impulsivity. Specifically, we leverage several electrophysiological, optical imaging and behavioral tools in combination with neuroeconomic and data-driven computational models to parse out aberrant patterns of neuronal encoding and dopamine release in response to natural and opioid reward cues. Importantly, we demonstrate that distinct endophenotypic motivational traits differentially underlie the enhanced propensity to excessively seek food and drug rewards and confirm that male subjects are at a greater risk of addiction-like vulnerability compared to their female counterparts.

## Results

### Prenatal THC exposure enhances effortful motivation for natural rewards in adult rats

To test if PCE augments motivation for rewards in the adult offspring, we administered the main psychoactive component of the *Cannabis sativa* plant THC (2 mg/kg, s.c., once daily) to rat dams from gestational day (GD) 5 until GD20 (Figure **1A**), a dosing regimen devoid of effects on litter size, maternal care, and offspring bodyweight^14^. The vehicle-(CTRL) and THC-exposed (PCE) progeny was left undisturbed until postnatal day (PND) 90, when surgical procedures took place. After surgical recovery, or ∼PND120, male and female rats where food-restricted and trained to lever press for palatable food rewards under various schedules of reinforcement (Figure **1B**). Results from the fixed ratio 1 (FR1) test indicate no differences due to PCE at low response requirements (Figure **1C**).

**Figure 1.**
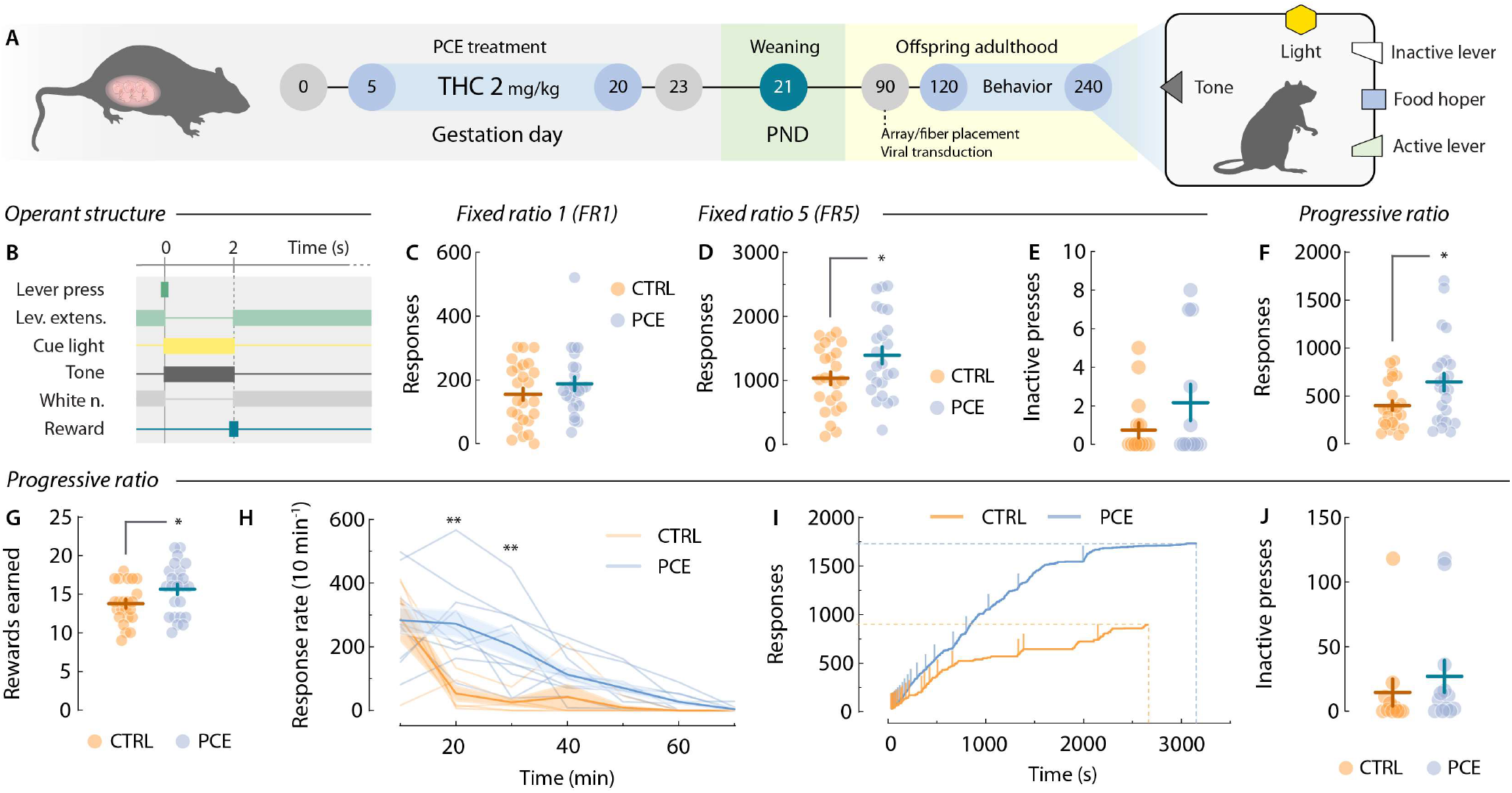
Motivation for food rewards in adult control and PCE progenies. **A**) Schematic timeline and experimental set-up of food self-administration experiments. **B**) Schematic representation of operant task structure of FR1, FR5 and PR schedules of reinforcement. **C**) Total active lever presses from the last FR1 training session (*t*_47_ = 1.12, *p* = 0.26). *n*_CTRL_ = 25; *n*_*PCE*_ = 24. **D**,**E**) Total active (*t*_45_ = 2.16, \**p* = 0.036) and inactive (Welch’s *t*_45_ = 0.95, *p* = 0.34) lever presses from the last FR5 operant session. *n*_CTRL_ = 23; *n*_PCE_ = 24. **F**,**G**) Total active lever presses (Welch’s *t*_35.8_ = 2.36, *p* = 0.023) and rewards earned (*t*_45_= 2.1, *p* = 0.034) under a PR schedule of reinforcement. *n*_CTRL_ = 23; *n*_PCE_ = 24. **H**) Average lever pressing frequency during PR responding (two-way RM ANOVA; ^*treatment*^*F*_1,13_ = 7.48, *p* = 0.017; ^*time*^*F*_6,78_ = 22.4, *p* < 0.001; ^*interaction*^*F*_6,78_ = 4.24, *p* < 0.001) (Bonferroni’s post-hoc; \*\**p* > 0.001). Thick lines and shaded areas represent mean ± SEM values. *n*_CTRL_ = 5; *n*_PCE_ = 10. **I**) Representative cumulative responses during PR. Vertical lines denote pellet deliveries. **J**) Total inactive PR lever presses (*t*_45_ = 0.74, *p* = 0.46). For all bar and point plots, lines represent mean ± SEM.

Rats then progressed to a FR5 schedule. Under these conditions, PCE offspring showed increased motivation (Figure **1D**), without changes in inactive lever pressing (Figure **1E**). Next, we tested rats on a progressive ratio (PR) schedule, where the response requirement grew exponentially to obtain a single palatable pellet. This measure of effortful motivation was significantly increased in terms of both active lever-pressing (Figure **1F**) and break-points (rewards earned) (Figure **1G**). Figure **1H** depicts average response rates during the PR session, demonstrating higher response vigor in PCE rats compared to control rats. For reference, Figure **1I** displays cumulative response traces from two representative rats during PR. No differences in inactive lever presses were found (Figure **1J**). Additionally, no significant effects of sex were found (Supplementary Figure **1**). Altogether, these results align with clinical findings suggesting a link between PCE and a propensity to manifest enhanced motivation later in life^11^. These findings highlight the utility of our rat model to study this clinically-relevant phenotype while circumventing residual confounding factors inherent to clinical studies.

### PCE results in a generalized enhancement of cue-evoked dopamine release in the NAc

Vigorous appetitive behaviors triggered by outcome-predictive cues are critically dependent on NAc dopamine release^20^. As such, striatal dopamine abnormalities underlie different motivational disorders, such as drug addiction, *via* dysregulated rewarding behaviors^21^. We have previously shown that PCE shifts the excitatory-to-inhibitory synaptic input balance onto ventral tegmental area (VTA) dopamine neurons^14^, resulting in increased excitability and firing rates^13^. However, whether the increase in reward seeking is accompanied by an enhancement of terminal dopamine release in a behaving animal is yet to be determined. Therefore, we used the fluorescent sensor GrabDA_2m_^22^ to monitor *in vivo* NAc dopamine fluctuations during the PR test.

Freely-moving rats implanted with optic fibers aimed at the core sub-field of the NAc were connected to a fiber photometry system via a patch chord relaying dopamine (470 nm)- and isosbestic (405 nm)-stimulated fluorescence (Figure **2A**). The raw 470 nm-stimulated dopamine signal exhibited fluctuations time-locked to completion of each PR response requirement (Figure **2B**). Post-mortem immunohistochemistry confirmed the localized expression of the GrabDA_2m_ probe and the optic fiber placement in the NAc core (Figure **2C**). Potential movement artifacts were subtracted by de-trending isosbestic fluctuations from the 407 nm-stimulated signal (Figure **2D**). Trial-by-trial traces from representative rats depict dopamine encoding of reward-paired cues (Figure **2E**). Importantly, PCE rats exhibited an increase in the area under the curve (AUC) associated with dopamine fluorescence during the 2-sec cue window, but not before (-2 to 0-sec) or after (2 to 4-sec) (Figure **2F**). Figure **2G** shows the trial-averaged peak GrabDA_2m_ amplitudes observed during cue presentation, revealing a significant increase of NAc dopamine release triggered by outcome-predictive cues.

**Figure 2.**
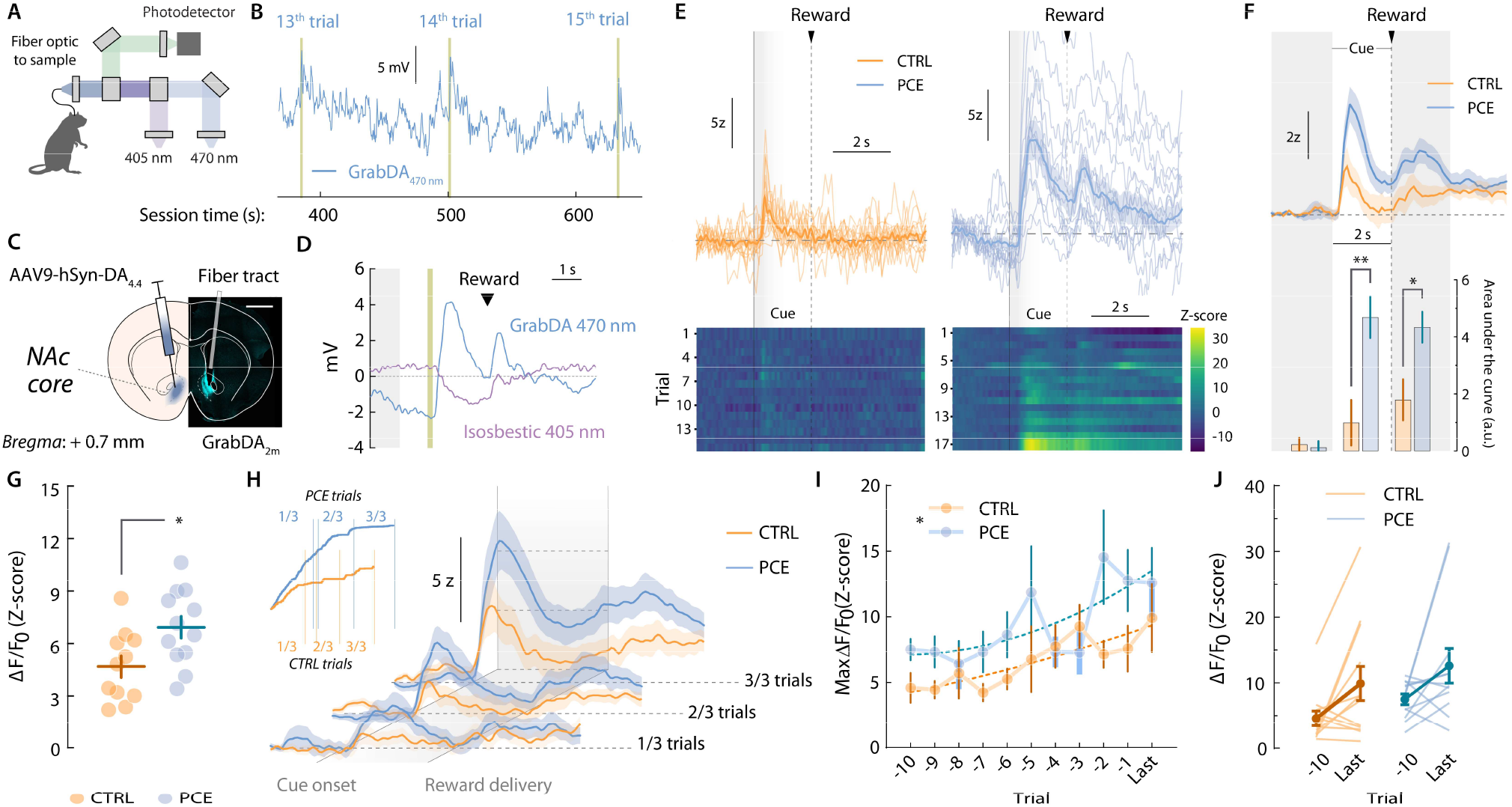
Cue- and reward-evoked NAc dopamine release events during food PR responding. **A**) Schematic representation of fiber photometry equipment consisting of dopamine-reporting 470 nm- and artifact-reporting 405 nm-stimulated light signals. **B**) Raw 470 nm-stimulated stream signal illustrating spontaneous and cue-aligned NAc dopamine transients during PR responding. Yellow bars represent cue presentation. **C**) Confocal and schematic image showing unilateral injection of the GrabDA_2m_ sensor and fiber tract aimed at the NAc core. Scale bar = 200 μm. **D**) Representative peri-stimulus time histogram (PSTH) depicting raw GrabDA_2m_ (470 nm) and isosbestic (405 nm) fluctuations time-locked to cue presentation (yellow bar) and reward delivery. Grey shaded area shows baseline period used for Z-scoring. **E**) Representative heatmap and trial-by-trial dopamine transients during PR responding. Grey shaded areas (0-2sec relative to cue onset) correspond to the time of cue presentation. **F**) (top) Group-averaged NAc dopamine transients during PR responding. (bottom) Pre-, cue- and reward-associated dopamine amplitudes (AUC) in control and PCE rats (two-way RM ANOVA; ^*treatment*^*F*_1,22_ = 5.60, *p* = 0.027; ^*event*^*F*_2,44_ = 17.12, *p* < 0.001; ^*interaction*^*F*_2,44_ = 5.09, *p* = 0.010 –followed by Bonferroni’s post-hoc; \*\**p* = 0.009). **G**) Cue-associated (0-2sec relative to cue onset) GrabDA_2m_ peak Z-score values (*t*_22_= 2.70, \**p* = 0.013) during PR. **H**) Group-averaged GrabDA_2m_ transients on each PR third. (inset) Representative cumulative responding plot illustrating separation of early (1/3), mid (2/3), and late (3/3) PR sections based on each animal’s total number of completed trials. **I**,**J**) Cue-associated peak GrabDA_2m_ (Z-scores) values from the last 10 completed PR trials (two-way RM ANOVA; ^*treatment*^*F*_1,22_ = 4.97, \**p* = 0.037; ^*trial*^*F*_10,220_ = 2.77, *p* = 0.003; ^*interaction*^*F*_10,220_ = 1.11, *p* = 0.35). Vertical lines represent SEM values. For all panels, *n*_CTRL_ = 12; *n*_PCE_ = 12.

Overall, NAc dopamine cue-evoked responses increased as a function of response requirement (Figure **2H**)^23^. Different reasons can account for this behavior, including the signaling of sunk costs^24,25^ and prediction errors^26^. To explore potential consequences of PCE in these phenomena, we compared the cue-evoked GrabDA_2m_ amplitudes of the last 10 ratios completed (Figure **2I, J**). The lack of interaction (^*interaction*^F_10,220_ = 1.11, *p* = 0.35) between trial (effort level) (^*trial*^F_10,220_ = 2.77, *p* = 0.003) and treatment (^*treat*^F_1,22_ = 4.97, *p* > 0.001) supports a baseline propensity for PCE offspring to show greater cue-evoked dopamine responses at all response requirement levels. This vulnerability was independent from potential dopamine fluctuations introduced by sunken cost valuation or error predictions.

### PCE enhances the encoding of cue-responding, effort-related NAc cell assemblies

Following these and prior dopaminergic findings^13,14^, we explored potential post-synaptic maladaptations. Using *in vivo* multiple single-unit electrophysiological recordings in control and PCE rats, we monitored the activity of NAc neurons during PR (Figure **3A**). Based on peri-event firing patterns (Figure **3B**), we performed a supervised *k*-means clustering analysis that identified four neuronal responses time-locked to trial completion (Figure **3C**). These clusters were characterized by firing rate increases or decreases at cue (purple, violet) or reward onsets (green, grey) (Figure **3D**) and did not vary as a function of the clustering algorithm applied (Supplementary Figure **2**).

First, we compared the relative abundance of encoding patterns on each treatment condition (Figure **3E**). Bayesian logistic regression models were created at each binary classification step (‘encoding vs. non-encoding’, ‘cue onset vs. reward onset’, ‘activated vs. inhibited’), and *posterior* distributions (β) were used to ascertain if the likelihood of finding a biased neuronal classification was higher to that of the null (control) model. Although PCE did not influence the proportion of stimuli-responsive NAc units (Figure **3F**), it biased them to preferentially encode reward-predictive cues instead of rewards (Figure **3G**). PCE, however, did not bias the direction (activation vs. inhibition) of the encoding (Figure **3H**). A closer examination revealed that cue-responding neurons from PCE rats fired at higher rates during cue presentation (0-2 sec) (Figure **3I**). There were no differences in the reward-activated (2-7 sec) (Figure **3J**) or cue-inhibited (0-2 sec) (Figure **3K**) clusters; however, the reward-inhibited cluster presented a larger rebound in activity after inhibition (Figure **3L**). These experiments expand NAc dopamine findings by disclosing an accompanying post-synaptic functional adaptation imposed by PCE, causing a disproportional representation of reward-paired cues during operant responding.

We next asked whether differential encoding dynamics between treatment groups dynamically diverged as a function of reward cost. To address this question, we employed a novel unsupervised algorithm to uncover demixed, low-dimensional neuronal dynamics across multiple timescales^27^. Briefly, we applied tensor component analysis (TCA), which extracts distinct cell assemblies (clusters) from large-scale recordings and groups them based on a three-dimensional low-rank tensor^28^ reflecting (*1*) the within-trial peri-event time histogram (PSTH) encoding pattern (*a*^*n*^_*p*_, Figure **3M**), (*2*) the between-trial temporal evolution of such encoding pattern (*b*^*n*^_*t*_, Figure 3N) and (*3*) the contribution of each recorded neuron to a given encoding pattern (*w*^*n*^_*b*_, Figure **3O**). Compared to photometry data, multiple single-unit recordings are better suited for TCA but, to normalize the scale of dimension *b*^*n*^_*t*_ across subjects, we had to restrict our dataset to the first eight trials completed during PR (Figure **3P**), which was the lowest break-point achieved on that session. This allowed us to probe how NAc encoding dynamics evolved across response requirements, encompassing reward costs from 1 to 20 lever presses per pellet. TCA discovered *n* = 4 tensorial cell assemblies, resembling those obtained by *k*-means clustering (Figure **3Q-T**). This approach, however, allowed us to determine that the cue-activated cell assembly (*a*^*1*^_*p*_*b*^*1*^_*t*_*w*^*1*^_*b*_) (Figure **3Q**, top panel) progressively became offline as response requirement increased throughout the PR session (Figure **3Q**, mid panel). Of note, this assembly was more prominent in units recorded from PCE offspring (Figure **3Q**, bottom panel). Figure **3R** displays the reward-activated cell assembly *a*^*2*^_*p*_*b*^*2*^_*t*_*w*^*2*^_*b*_ (top panel), which remained constant across response requirements (middle panel) and treatment groups (bottom panel). The cue-inhibited tensor *a*^*3*^_*p*_*b*^*3*^_*t*_*w*^*3*^_*b*_ (Figure **3S**, top panel), however, progressively became online as effort increased (Figure **3S**, middle panel) and was again more evident in units recorded from PCE rats (Figure **3S**, bottom panel). Lastly, the reward-inhibited cell assembly *a*^*4*^_*p*_*b*^*4*^_*t*_*w*^*4*^_*b*_(Figure **3T**, top panel) did not vary throughout the PR session (Figure **3T**, mid panel) and exhibited similar loadings from control and PCE units (Panel **3T**, bottom panel). The effects of PCE present in cell assembly loadings were not modulated by sex (Supplementary Figure **3**).

**Figure 3.**
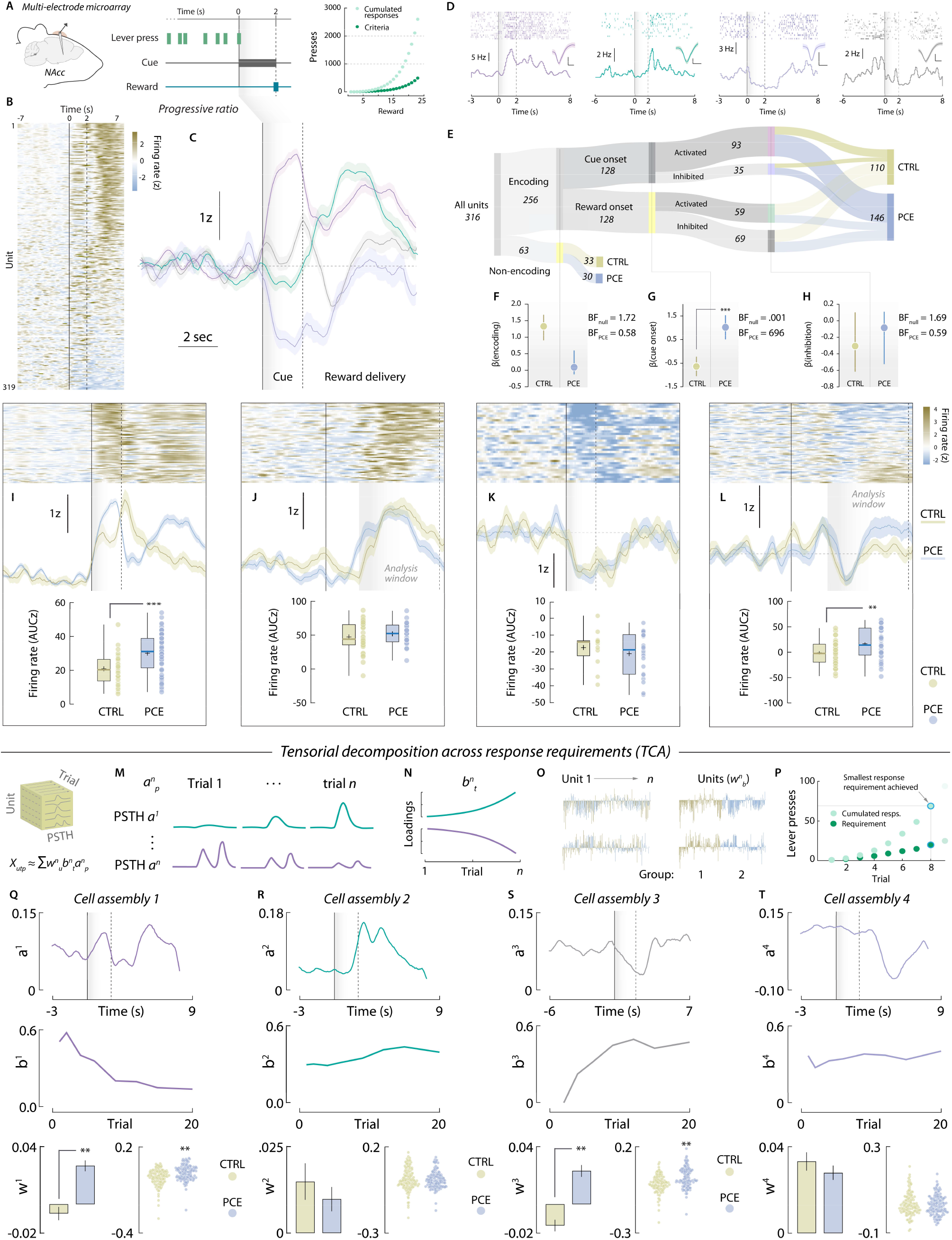
PCE promotes NAc cue-evoked neuronal firing in a temporally engendered manner. **A**) Schematic representation of chronic multi-electrode microarray NAc implantation and PR operant task structure. The right panel shows response requirements of each PR trail and its associated cumulated responses. **B**) Trial-averaged, cue-aligned PSTH of all recorded NAc units during PR responding. **C**) Time-course of the four distinct NAc firing rate patterns uncovered by *k*-means clustering. **D**) Raster (top) and PSTH (bottom) plots of representative neurons pertaining to each cluster. Insets represent average waveforms (scale bar: 50 μV, 20 ms). **E**) Sankey plot showing absolute amount (inset numbers) of encoding vs. non-encoding, cue onset vs. reward onset, and activated vs. inhibited units and its treatment group assignment. **F-H**) Posterior coefficients (β) with credible intervals (vertical lines) of Bayesian logistic regression models evaluating the influence of PCE on each classification step. Bayes factors (BF) for null and PCE-influenced hypotheses are shown as insets. ‘***’ Represents ‘extreme’ evidence in favor of the alternative (PCE-influenced) hypothesis. **I-L**) (Top) Trial-averaged heatmap depicting all neurons’ activity from each cluster. (Mid) Group-averaged NAc firing rate (Z-score) centered around cue onset (vertical black line). Shaded grey areas and vertical dashed lines indicate the time window used for parametric analyses. Lines represent average firing rates and colored shaded areas depict ± SEM. (Bottom) Trial-averaged firing rates (AUC) during analysis window from the cue-activated (*t*_91_= 3.64, \*\*\**p* > 0.001) (*n*_CTRL_ = 25; *n*_PCE_ = 68), reward-activated (*t*_57_= 0.88, *p* = 0.38) (*n*_CTRL_ = 35; *n*_PCE_ = 29), cue-inhibited (*t*_33_= 0.85, *p* = 0.39) (*n*_CTRL_ = 13; *n*_PCE_ = 22) and reward-inhibited units (\*\**t*_67_= 2.72, *p* = 0.008) (*n*_CTRL_ = 37 ; *n*_PCE_ = 32). For box plots, the center line represents the median; the cross illustrates the average; the bounds of the box depict the 25^th^ to 75^th^ percentile interval; and the whiskers represent the minima and maxima. **M-O**) Tensor representation of trial-structured neural data used to parse out effort-related NAc encoding dynamics during PR responding. **O**) Example unit factor loadings color-coded by group assignment. In the right panel, the same units have been ordered by group assignment. **P**) PR trials, associated lever presses and response requirements selected for TCA analysis. **Q-T**) PSTH pattern (top), trial-by-trial temporal evolution (mid) and unit factor loadings (bottom) of the four tensorial cell assemblies identified by TCA. PCE selectively upregulates effort-related, cue-triggered patterns of neuronal activity in the NAc, as reported by cell assemblies *a*^*1*^_*p*_*b*^*1*^_*t*_*w*^*1*^ (\*\**t*_317_ = 5.62, *p* < 0.001) and *a*^*3*^_*p*_*b*^*3*^_*t*_*w*^*3*^_*b*_ (\*\**t*_317_ = 6.41, *p* < 0.001), but not *a*^*2*^_*p*_*b*^*2*^_*t*_*w*^*2*^_*b*_ (Welch’s *t*_254.5_ = 0.79, *p* = 0.42) and *a*^*4*^_*p*_ *b*^*4*^_*t*_*w*^*4*^_*b*_ (*t*_317_= 0.98, *p* = 0.32) (*n*_CTRL_ = 143; *n* _PCE_ = 176).

Of note, we asked whether the four tensorial cell assemblies were functionally implicated in the effortful pursuit of food rewards. To do this, we trained a machine learning algorithm (gradient boosted decision trees) to predict the latency to obtain the 8^th^ PR reward –a behavioral metric not included in the TCA algorithm up until this point. The algorithm was trained on each unit’s loadings to its four tensorial assemblies (*w*^*1-4*^_*b*_) and significantly predicted the latency to obtain the 8^th^ reward (*ground truth* vs. *predicted value*; Pearson’s *r* = 0.44, *p* = 0.003) (Supplementary Figure **4**).

Altogether, these results uncover an additional source of alterations in NAc neuronal dynamics after *in utero* exposure to THC, linking the higher propensity to work harder with an overreactive representation of cues as effort increases. Intriguingly, this dynamic effect contrasts with the monolithic profile of PCE on NAc dopamine release, which did not vary according to reward cost. The combination of these effects points to additional PCE disturbances –beyond or downstream dopamine– in ventral striatal integrative processes regulating effortful motivation across declining rates of reinforcement.

### PCE increases voluntary opioid drug-seeking and relapse in a sex-dependent manner

Clinical cohorts and longitudinal studies point to an increased risk of substance use in adolescent and young adults prenatally exposed to cannabis^18,19,29^. This phenomenon has been replicated in PCE rodents, which exhibit marked behavioral adaptations when exposed to drugs of abuse later in life^30–33^. Notably, the risk of subsequent drug use (in this case, cannabis) was higher for male individuals relative to their female counterparts^18^. This agrees with the male-specific hyperdopaminergic findings described in male PCE rats^14^. With this evidence at hand, as well as the *in vivo* dopaminergic and food reward processing disturbances described here, we determined whether effortful motivation for a drug (opioid) reward was also enhanced, and whether potential sex differences could arise when pursuing drug reinforcers. Control (n = 27) and PCE (n = 22) male and female adult rats were subjected to jugular vein catheterization surgery to allow for voluntary i.v. infusions of the fast-acting fentanyl analog remifentanyl (1 µg/kg/inf) (Figure **4A**). We chose remifentanyl for its short half-life, which clears from blood in ∼0.3 min and ∼10 min from the NAc^34^. Its rapid clearance reduces the risk of overdose, rendering it ideal for rodent intravenous self-administration (IVSA) studies^35^ and within-session behavioral economics tests (see below) ^36^. This, in combination with its abuse potential in humans^37^, makes remifentanyl a safe, convenient, and relevant tool for rodent studies on opioid use.

**Figure 4.**
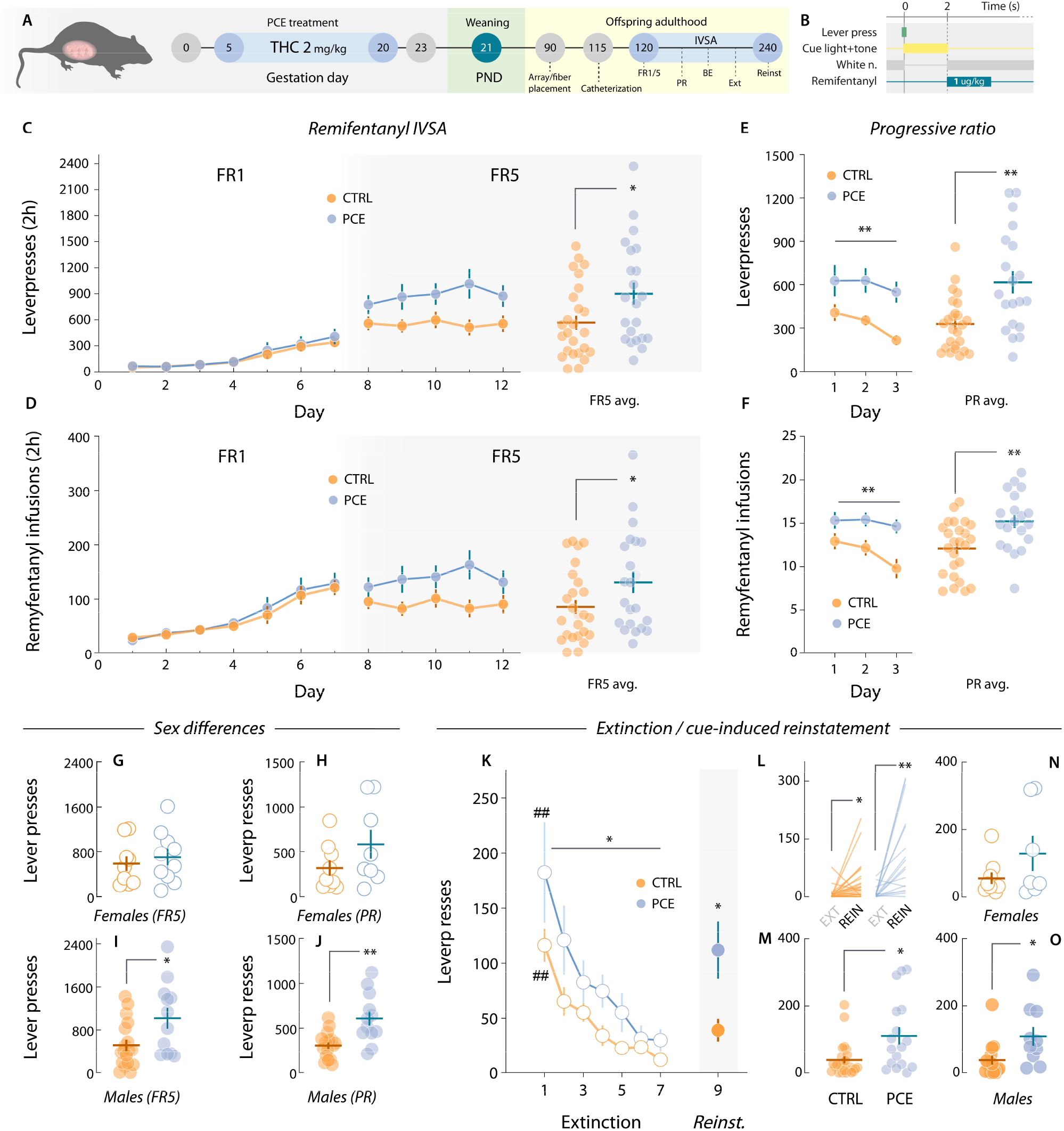
Motivation for drug (remifentanyl) rewards in adult PCE rats. **A**) Schematic timeline of PCE treatment and remifentanyl i.v. self-administration experiments. **B**) Remifentanyl operant tasks structure. **C**) Daily active lever presses under FR1 (two-way RM ANOVA; ^*treatment*^*F*_1,47_ = 0.19, *p* = 0.66; ^*day*^*F*_6,265_ = 20.7, *p* < 0.001; ^*interaction*^*F*_6,265_ = 0.44, *p* = 0.96) and FR5 schedules of reinforcement (two-way RM ANOVA; ^*treatment*^*F*_1,47_ = 5.43, *p* = 0.024; ^*day*^*F*_4,188_ = 1.09, *p* = 0.35; ^*interaction*^*F*_4,188_ = 1.98, *p* = 0.09). The right dot plot shows average FR5 values (\**t*_47_ = 2.39, *p* = 0.024). *n*_CTRL_ = 27; *n*_PCE_ = 22. **D**) Daily remifentanyl infusions obtained under FR1 (two-way RM ANOVA; ^*treatment*^*F*_1,47_ = 0.07, *p* = 0.77; ^*day*^*F*_6,262_ = 25.5, *p* < 0.001; ^*interaction*^*F*_6,262_ = 0.53, *p* = 0.77) and FR5 (two-way RM ANOVA; ^*treatment*^*F*_1,47_ = 4.11, *p* = 0.048; ^*day*^*F*_4,188_ = 1.07, *p* = 0.37; ^*interaction*^*F*_4,188_ = 2.14, *p* = 0.07). The right dot plot shows average FR5 infusions (\**t*_47_ = 2.06, *p* = 0.048). *n*_CTRL_ = 27; *n*_PCE_ = 22. **E**) Daily (two-way RM ANOVA; ^*treatment*^*F*_1,43_ = 12.5, \*\**p* = 0.001; ^*day*^*F*_2,86_ = 4.95, *p* = 0.009; ^*interaction*^*F*_2,86_ = 0.54, *p* = 0.58) and average (Welch’s \*\**t*_27.9_ = 3.35, *p* = 0.002) lever presses during PR testing. **F**) Daily (two-way RM ANOVA; ^*treatment*^*F*_1,39_ = 8.11, \*\**p* = 0.007; ^*day*^*F*_2,78_ = 6.04, *p* = 0.003; ^*interaction*^*F*_2,78_ = 3.83, *p* = 0.025) and average (\*\**t*_39_ = 2.85, *p* = 0.007) PR remifentanyl rewards. *n*_CTRL_ = 25; *n*_PCE_ = 20. **G**,**H**) Female rat lever pressing levels during FR5 (*t*_17_ = 0.58, Bonferroni-Dunn-corrected *p* = 0.99) (females, *n*_CTRL_ = 9; *n*_PCE_ = 10) and PR sessions (*t*_15_ = 1.50, Bonferroni-Dunn-corrected *p* = 0.30) (females, *n*_CTRL_ = 9; *n*_PCE_ = 8). **I**,**J**) Average male lever pressing levels during FR5 (\*\**t*_26_ = 3.11, Bonferroni-Dunn-corrected *p* = 0.008) (males, *n*_CTRL_ = 18; *n*_PCE_ = 12) and PR (\*\**t*_24_ = 3.87, Bonferroni-Dunn-corrected *p* = 0.001) (males, *n*_CTRL_ = 16; *n*_PCE_ = 12).

As with food rewards, remifentanyl infusions (2-s) were preceded by a 2-s multi-sensorial compound cue (light + tone) (Figure **4B**). Both control and PCE rats learned to lever press at similar rates (Figure **4C**). On FR5, PCE rats started to exhibit increased levels of lever pressing (Figure **4C**).

Figure **4D** shows drug infusions obtained throughout FR1 and FR5, also significantly different. Next, rats underwent three IVSA PR sessions, in which PCE rats displayed higher levels of effortful motivation to work for remifentanyl in terms of total lever presses (Figure **4E**) and drug infusions (break-points) (Figure **4F**). Potential sex differences in these metrics were explored by comparing sex-matched subjects across treatments. No significant differences in FR5 (Figure **4G**) and PR lever pressing (Figure **4H**) were found among females. However, in accordance with clinical evidence^18^, male PCE offspring displayed higher levels of remifentanyl seeking during FR5 (Figure **4I**) and PR sessions (Figure **4J**). Overall, results confirm that the hypermotivational phenotype associated with PCE extends to drug rewards, –in particular the synthetic opioid remifentanyl–. Interestingly, unaffected FR1 performance suggests, once again, that no gross learning effects could explain the observed changes in motivation. Unlike with natural rewards, the consequences of PCE in remifentanyl-seeking rats were influenced by sex, conferring a higher vulnerability to the male offspring, and recapitulating the clinical phenotype.

Persisting risk of relapse remains a defining feature of substance use disorders and the primary endpoint for treatment considerations^38,39^. Owing to the increase in opioid intake and seeking uncovered here, we next aimed to parse out PCE effects on remifentanyl seeking in drug-free conditions (extinction) and upon re-exposure to drug-paired cues (reinstatement), a rodent model of relapse^40^. PCE rats showed increased overall drug seeking levels during extinction, but the rate at which they ceased seeking the drug-paired lever was not different (Figure **4K**). Upon re-introduction of drug cues, both control and PCE rats increased their respective levels of opioid-seeking compared to the last extinction session (Figure **4L**). Notably, PCE rats exhibited greater levels of cue-induced reinstatement (Figure **4L,M**). We did not detect this increase when comparing females across treatments (Figure **4N**). However, male PCE offspring showed a marked increase in cue-induced reinstatement when compared to male control rats (Figure **4O**).

### PCE exacerbates opioid-induced dopamine release in the NAc of PCE animals

The reinforcing potential of abuse drugs depends on its ability to increase the concentration of dopamine in certain brain structures^41,42^, such as the ventral striatum^43^. To probe drug-evoked dopaminergic alterations imposed by PCE, we implanted optic fibers and the GrabDA_2m_ sensor in the NAc of remifentanyl-seeking rats (Figure **5A**).

‘Frist, drug-naïve, catheterized rats were left undisturbed in an operant chamber (no manipulanda available) for 10 min before receiving a non-contingent high-dose remifentanyl infusion (10 μg/kg). Spontaneous NAc GrabDA_2m_ fluctuations were recorded for 15 min and dopamine release events were identified based on their prominence (Methods) (Figure **5B, C**). Pre- and post-drug release events were distributed based on their amplitudes (Figure **5B,C** insets). Results indicate that, while both groups exhibited larger-amplitude NAc dopamine release events post-drug infusion, PCE amplified this response (Figure **5D,E**), revealing a neuropharmacological propensity to experience larger dopaminergic responses to remifentanyl.

We then assessed if PCE enhances the encoding of drug-paired cues during voluntary pursuit of remifentanyl. Control and PCE animals were recorded during PR responding. Trial-by-trial representative recordings reveal phasic GrabDA_2m_ transients time-locked to the onset of the drug-paired cues (Figure **5F**). Group-averaged measurements confirm this pattern of encoding (Figure **5G**), but do not disclose differences in the detected peak amplitudes during the cue (0-2s) and drug infusion (2-8s) time windows (Figure **5H**). In spite of this, further analysis revealed sex-specific differences. Accordingly, no differences in cue- or drug-evoked dopamine encoding were found among female control and PCE rats (Figure **5I,J**). Notably, a specific increase in dopaminergic encoding of opioid-paired cues was found among PCE males (Figure **5K,L**). Of note, the male-specific enhancement in opioid cue encoding agrees with the previously disclosed PCE male willingness to work harder for remifentanyl.

### Male PCE offspring NAc neurons disproportionately signal drug-paired cues

We next evaluated if ventral striatal neuronal firing dynamics were similarly disturbed. In this case, a subset of control and PCE rats underwent chronic multi-electrode implantation surgery (Figure **6A,B**) and their NAc neuronal responses were monitored during voluntary pursuit of remifentanyl (PR) (Figure **6C**). The same *k*-means clustering algorithm used on the food PR dataset was applied, this time identifying three distinct modes of firing in response to trial completion (Figure **6D**). We found that NAc units either preferentially increased their firing rates during cue presentation (0-2s relative to cue onset) or remifentanyl infusion (2-7s relative to cue onset). A third, more modest subpopulation started inhibiting its firing rates prior to the preceding lever press. To detect potential encoding biases introduced by PCE, we performed a Bayesian logistic regression model comparing the relative proportion of NAc units pertaining to the cue-responding cluster. The likelihood of finding a PCE unit in the cue-responding cluster was higher than that observed for control units (Figure **6D** inset), confirming that drug-paired cues become over-represented in functionally-defined subpopulations of the NAc.

We next focused our attention on the cue-responding subpopulation (Figure **6E,F**). To compare the relative strength of encoding within this cell assembly, we obtained normalized firing rates (AUC) during the cue presentation time window (0-2s relative to cue onset) and found no gross differences between control and PCE rats (Figure **6G**). However, when we segregated firing rate data by sex to look for a potential male vulnerability, we found that female PCE offspring neurons fire at similar rates than their sex-matched control counterparts (Figure **6H**). Importantly, cue-responding NAc neurons of male PCE rats fired at considerably higher rates (Figure **6I**), further supporting the idea that adult male PCE rats are especially vulnerable to the reinforcing, dopaminergic and striatal actions of remifentanyl. No differences were found on the other two clusters (Supplementary Figure **5**).

### Distinct reward processing endophenotypes underlie PCE-induced excessive motivation

Increased reward-directed behaviors for food and remifentanyl (*1*), augmented cue-evoked dopamine release (*2*), and subsequent effort-related NAc neuronal dynamics (*3*) all accompany a motivational endophenotype likely predisposing PCE individuals to drug addiction. Nonetheless, these three outcomes can also derive from a more parsimonious, non-pathological neuropsychological adaptation. That is, should PCE animals experience greater hedonic responses to appetitive rewards, adaptations *1*-*3* would be interpreted as adaptive, as they would ensure the consecution of higher-valued goods. Resolving this conundrum is crucial to avoid mistaking a vulnerability to drug seeking for a more innocuous hedonic value-seeking phenotype.

We sought to resolve this ambiguity by exploiting neuroeconomic modelling tools on an exponential demand task^44^. After PR testing, food-seeking control and PCE rats progressed to the demand task. During five 50-min long sessions, response requirements (or ‘price’) progressively increased every 10-min and averaged demand levels (pellets obtained) per price point were fitted using a neuroeconomic exponential demand equation^45^ (Figure **7A**). The parameters *Q*_*0*_, *α, P*_*max*_, and *EV* were then used to map distinct reward valuation processes as a function of PCE. *Q*_*0*_ is denoted as the reward quantity consumed at zero price (Figure **7A**, *y*-intercept), and corresponds to the reward’s hedonic set point^46,47^. Control and PCE animals shared *Q*_*0*_ levels (Figure **7B**). Unaffected *Q*_*0*_ values indicate that the palatability of food pellets was the same for PCE rats and, therefore, the hypermotivational and hyperdopaminergic phenotypes described must derive from other sources of reward valuation. One alternative is price sensitivity^48^, which is denoted by *α* and is orthogonal to *Q*_*0*_. In this case, increased effortful motivation would be explained by decreased sensitivity (*α*) to price increments (i.e., response requirements), making rats insensitive to increasing difficulty and displaying greater break-points. However, *α* values of PCE rats were equivalent to those of control subjects (Figure **7C**). A third possibility is an increase in the inflection point between inelastic and elastic demand, represented by *P*_*max*_. *P*_*max*_ is related to *α* and indicates the threshold value (in response requirement) at which price sensitivity sets in. In other words, *P*_*max*_ coincides with the peak of responding rates and is interpreted as motivational drive; that is, the maximum amount of behavioral resources a subject is willing to allocate to obtain a standard-valued reward^47^. We found that PCE animals exhibited much larger *P*_*max*_ values (Figure **7D**), thereby supporting an excessive motivational drive that is independent of hedonic set points (*Q*_*0*_) and sensitivity to declining opportunity costs (*α*). Figure **7E** shows that the essential value of food pellets –its reinforcing efficiency at all price points– was unaffected. Finally, Figure **7F** contains the raw demand task lever pressing data from which all parameters were derived.

Next, we performed a dimensionality reduction in all behavioral measurements obtained (all neuroeconomic parameters, FR1-5, and PR) to gain access to a more generic manifestation of the underlying motivational endophenotype^49^. Principal component analysis (PCA) identified two low-dimensional factors explaining 84.7 % of all behavioral variance (Figure **7G**). The first principal component (PC1) reflected inelastic lever pressing, as all variables related to lever pressing (*P*_*max*_, *Q*_*0*_, FR5 responses, PR break-points) loaded positively onto this dimension, while elasticity (*α*) loaded negatively (Figure **7H**). Both control and PCE rats scored similarly in this dimension (Figure **7I**), confirming that this food-seeking behavioral signature was not impacted. A second dimension, PC2, positively related *P*_*max*_ and PR break-points with small *Q*_*0*_ (Figure J), indicative of “anhedonic” motivational drive. It is this endophenotype that clearly characterized the motivational disturbances regarding natural rewards, as THC-exposed rats scored much higher in this dimension (Figure **7K**). Altogether, these results confirm that adult PCE offspring suffer from excessive motivation for natural rewards, even when these are less valued and in absence of benefit-cost calculations.

Because of their pharmacological effects^20^ and the lack of homeostatic compensatory mechanisms ^50^, drugs of abuse function differently compared to natural rewards (i.e., food pellets). Therefore, in order to investigate further signs of maladaptive behavior related to addiction-like behaviors, we conducted the same endophenotypic analysis on remifentanyl self-administering rats. Following PR testing, and prior to extinction sessions, control and PCE rats conducted five 80-min long remifentanyl exponential demand tasks (Figure **7L**). As with natural rewards, PCE augmented maximum motivational drive (*P*_*max*_) for remifentanyl but did not influence its related hedonic set point (*Q*_*0*_) or price sensitivity (*α*) (Figure **7M-O**). In addition, remifentanyl had a higher essential value (Figure **7P**). The essential value of a reward reflects its reinforcing efficiency at all price points and can be clearly visualized in the generalized increase in lever pressing observed during the demand task (Figure **7Q**). Principal component analysis revealed two low-dimensional signatures (Figure **7R-V**), resembling those of food seeking behavior. Intriguingly, when working for a drug reward, PCE induced a specific increase in inelastic lever pressing (PC1) (Figure **7T**) but not “anhedonic” motivational drive (PC2) (Figure **7V**). Altogether, these results demonstrate that distinct endophenotypic alterations underlying seemingly equivalent increases in natural and drug reward seeking behaviors are a hallmark of PCE.

### PCE rats exhibit higher impulsivity

Impulsivity deficits are one of the outcomes most consistently associated with PCE^51–55^ and are hypothesized to underlie increased risk of substance use observed after PCE^11^. Thus, a failure in inhibitory control required to interrupt or suppress highly-automatized reward-seeking behaviors could contribute to the PCE hypermotivational phenotype^56^. Hence, to examine whether PCE rats also presented heightened impulsive actions, we trained rats on a go/no-go task^57^. Go trials were signaled by a 5-sec compound cue (light + tone) and a lever press within cue presentation earned a palatable food pellet reward (Figure **8A**). No-go trials were signaled by a different 5-sec compound cue (house light + white noise) and were rewarded only if no lever press was detected during cue presentation (Figure **8B**). Control and PCE rats were trained on this task during 13 sessions (Figure **8C**). As all rats flawlessly executed nearly all go trials (Figure **8D**), the number of successfully executed no-go trials per session was considered a measure of each animal’s inhibitory self-control. Data from the final session, when impulsivity levels were overall lower, revealed that PCE rats displayed higher levels of impulsivity (Figure 8 **E,F**). In line with the male-specific PCE impairments examined so far, no significant differences in no-go performance were present among control or PCE female subjects (Figure **8G**). However, impulsivity differences reached statistical significance when control males were compared against PCE male subjects (Figure **8H**). Finally, a subset of rats progressed to a second phase of higher difficulty, in which lever pressing had to be withheld for 10-sec (Figure **8I**). A significant difference between control and PCE rats was also found at this level of inhibitory control (Figure **8J**). Therefore, we reveal another maladaptive trait associated to PCE and related to the ability to control highly-automatized reward-oriented motor actions^58^.

## Discussion

In the present study, we provide evidence that prenatal exposure to THC leads to modifications in striatum-based processing consistent with an increased risk for opioid reinforcement and, ultimately, relapse. These alterations encompass an increased willingness to work for natural and drug rewards, upregulated dopamine release responses to outcome-predictive cues, overrepresentation of effort-encoding NAc neural dynamics and elevated impulsivity. Collectively, these consequences of PCE correspond to well-established neurodevelopmental risk factors for drug use disorders^59,60^.

Individual differences contribute to the development of long-term maladaptations driving compulsive drug use and relapse propensity^61^. Maternal exposure to THC is such a predictive risk factor^13^. Nonetheless, little is known about the neuropsychological constructs leading to this propensity. In our study, we present neuroeconomical evidence of a detrimental trait influencing how PCE individuals work for natural and drug reinforcers. On one hand, the increased willingness of PCE offspring to work impulsively for natural (food) rewards is associated with their intrinsically higher motivational drive, even when accounting for individual differences in reward valuation. On the other hand, increased remifentanyl-seeking is a manifestation of a greater susceptibility to the reinforcing actions of opioids, independent of their hedonic effects. In terms of Berridge and Robinson’s (2016)^62^ framework of addiction^63^, these observations support the hypothesis of a hypersensitized ‘wanting’ system, in contrast to reward valuation (‘liking’) or learning alterations, which we (and others^17^) demonstrate are not implicated in the motivational phenotype of PCE. A specific increase in incentive salience provides a satisfactory explanation for this and prior rodent studies reporting enhanced effortful motivation and psychomotor activation^64^ in response to heroin^65,66^, food^17^, THC^14^ and alcohol^67^ (reviewed in ^11^).

Preclinical and clinical evidence have established a causative role for enhanced mesostriatal dopamine function in the attribution of incentive salience and, ultimately, the pathogenesis of addiction^68,69^. Accordingly, we report that augmented NAc dopamine release is a hallmark of PCE. This finding expands previous evidence indicating that VTA dopamine cells become hyperexcitable following maternal THC exposure^14^, and other reports showing increased drug-evoked dopamine synthesis^15^. In addition, our photometry read-outs reveal a specific upregulation of dopamine release in response to reward-paired cues, a dysregulation known to predispose subjects to compulsively seek drugs of abuse^70,71^ and a higher risk of relapse^20^. Specifically, reintroduction of opioid-paired cues triggered greater remifentanyl-seeking. Related to this, PCE rats exhibited signatures of hyper-responsive NAc dopamine activity, even in drug-naïve conditions. It has been proposed that decreased expression of NAc dopamine D2 receptors in human PCE offspring may contribute to such potentiated state^72^. Future studies should examine whether this pharmacodynamic susceptibility, possibly related to impaired D2 auto-receptor function, can explain the specific increase in opioid reinforcing efficiency seen after maternal THC.

Dopamine release in the NAc gates striato-thalamic circuits, coupling internal states, exteroceptive stimuli and motor invigoration mechanisms to ensure proper behavioral tunning to constantly-evolving environments^73,74^. Therefore, we asked whether the hyperdopaminergic and motivational disturbances associated with PCE were accompanied by changes in NAc neuronal patterns of encoding. We found that electrophysiological measurements matched dopamine findings, and a large bias towards outcome-predictive cues was determined. To explore the functional relationship of such neurobiological correlates with the behavioral phenotype observed, we opted for a machine learning-based approach. We decomposed the influence of PCE on NAc tensorial cell assemblies across declining rates of reinforcement^27^, linking response vigor progression during PR with temporal encoding dynamics of outcome-predictive cues. Moreover, we observe that PCE progeny NAc neurons are more aligned with cue-triggered, effort-engendered patterns. Therefore, dopaminergic and striatal processing alterations identified in this study are significant contributors of the hypermotivational susceptibility conferred by *in utero* THC exposure. Notwithstanding, it must be noted that not all consequences of PCE on NAc dopamine release and neuronal encoding must derive from a unique, common alteration of VTA dopamine cell hyperexcitability^12^. Unlike NAc encoding patterns, PCE effects on dopamine release were not modulated by decreasing rates of opportunity costs. This points to additional, overlapping ventral striatal modifications subsequent to PCE, which expand others described in the prefrontal cortex^75,76^, and hippocampus^77^.

Much like seen in human cohorts^11^, the risk for augmented drug reinforcement was higher for males. Of note, sex-dependent behavioral effects of PCE paralleled its neurobiological signatures. That is, in those instances in which the behavioral consequenceswere sex-specific (remifentanyl break-points), dopamine release and cue-evoked neuronal firing rates were also found exclusively upregulated in males. This phenomenon is consistent with sex-specific shifts in excitatory-to-inhibitory synaptic balance observed in VTA dopamine cells^14^, as well as NAc epigenetic alterations observed following PCE in males but not females^17^. Although we present an interpretative framework for addiction-related vulnerabilities, the causative mechanisms linking THC-triggered molecular cascades in the developing brain to the enduring neurobiological signatures identified here remain largely unknown. The comprehensive study of Tortoriello et al.^78^, however, suggests that fetal THC destabilizes microtubule organization in developing neurons and dampens CB1R influence in the assembly of fetal brain circuits^79^. It remains to be seen how fetal sex could interact with the affected substrates in order to promote male-specific vulnerability observed across neurobehavioral domains and species.

In conclusion, the persistent striatal processing and behavioral modifications identified here highlight a notable public health concern. The spurious neurodevelopmental effects of PCE accompany the propensity for late-onset motivational disturbances and potentially drug addiction issues, underscoring an urgent need for enhanced awareness and mitigation strategies regarding the use of cannabis during pregnancy.

## Acknowledgements

This research was supported by the National Institute on Drug Abuse (K99 DA060209 to M.A.L. and R01 DA022340 to J.F.C.

## Author contributions

Conceptualization: M.A.L. and J.F.C. Methodology: M.A.L., R.Y.M., S.A. and J.F.C. Formal analyses: M.A.L. Investigation: M.A.L. and R.Y.M. Data curation: M.A.L. and R.Y.M. Writing - original draft: M.A.L. Writing - review and editing: M.A.L. and J.F.C. Visualization: M.A.L. Supervision: J.F.C. and M.M. Project administration: J.F.C. Funding acquisition: J.F.C., I.K. and M.M.

## Materials and Methods

All experimental procedures conformed to the National Institute of Health *Guide for the Care and Use of Laboratory Animals*. Ethical approval was granted by the Institutional Animal Use and Care Committee at the University of Maryland, Baltimore (IACUC protocol #00000054).

## Resource availability

### Lead contact

Further information and requests for resources and reagents should be directed to and will be fulfilled by the lead contact, Joseph F. Cheer.

### Materials availability

The study did not generate any new unique reagents or materials. The materials and reagents employed herein can be accessed commercially without availability restrictions.

### Data and code availability

Access to the original data will be provided upon email request to the lead contact. The original code for *k*-means clustering will be made publicly available upon completion of peer revision. Any additional information required to reanalyze the data reported in this paper is available from the lead contact upon request.

## Experimental model and subject details

### Animals

For all experiments, male and female Sprague-Dawley rats were used. Animals were housed in a temperature- and humidity-controlled room (24 °C and 40–50% humidity, respectively) and maintained on a 12 h light/dark cycle (07:00–19:00h). All experiments were conducted in the light cycle. The number of rats for each experiment are indicated in the respective figure legends. Sex of the animals is reported throughout the figure legends. For all experimental measures, subjects from at least four different THC-exposed liters were included. All experimenters were blind to treatment assignments once liters were weaned at PND21.

## Method details

### Prenatal THC exposure

Primiparous female Sprague-Dawley rats (Jackson Laboratories), mated with males, were used as maternal subjects, and housed individually during pregnancy. Each day, from GD5 to GD20, they were administered either THC (2 mg/kg, 2 ml/kg, s.c.) or vehicle. THC (dissolved in a 100 mg/ml ethanol solution) was obtained from NIDA Drug Supply Program (NDSP). Then, it was emulsified in 2% Tween 80, sonicated for 5 min and dissolved in sterile physiological saline. This specific dose was selected because it has been shown to not induce the typical behavioral responses observed in the cannabinoid tetrad assay, nor does it lead to cannabinoid tolerance^80^. Previous studies by our group have demonstrated that this dose does not significantly affect maternal behavior, behavioral non-maternal activity, or the bodyweight of the offspring ^13,14^. This dose is also comparable to the THC concentration found in mild cannabis joints (5%) and reflects moderate human cannabis consumption levels ^81^.

Offspring were weaned at ∼PND21 and maintained without any further manipulation with access to food and water until experimental day (PND90). The study’s overall design was as follows: two independent cohorts of control and PCE subjects were used, one subjected to food-seeking operant training and the other to remifentanyl self-administration. In each cohort, a portion of the subjects underwent fiber photometry surgery, another portion were implanted with multiple-electrode arrays and a third portion did not undergo any NAc implantation procedure. No differences based on NAc implantation strategy were observed in terms of operant food- or remifentanyl-seeking. Figures 1 and 5 comprise behavioral measurements from all implanted and non-implanted rats. Data from the exponential demand (Figure **7**) and go/no-go tasks (Figure **8**) were obtained from the non-implanted set rats.

**Figure 5.**
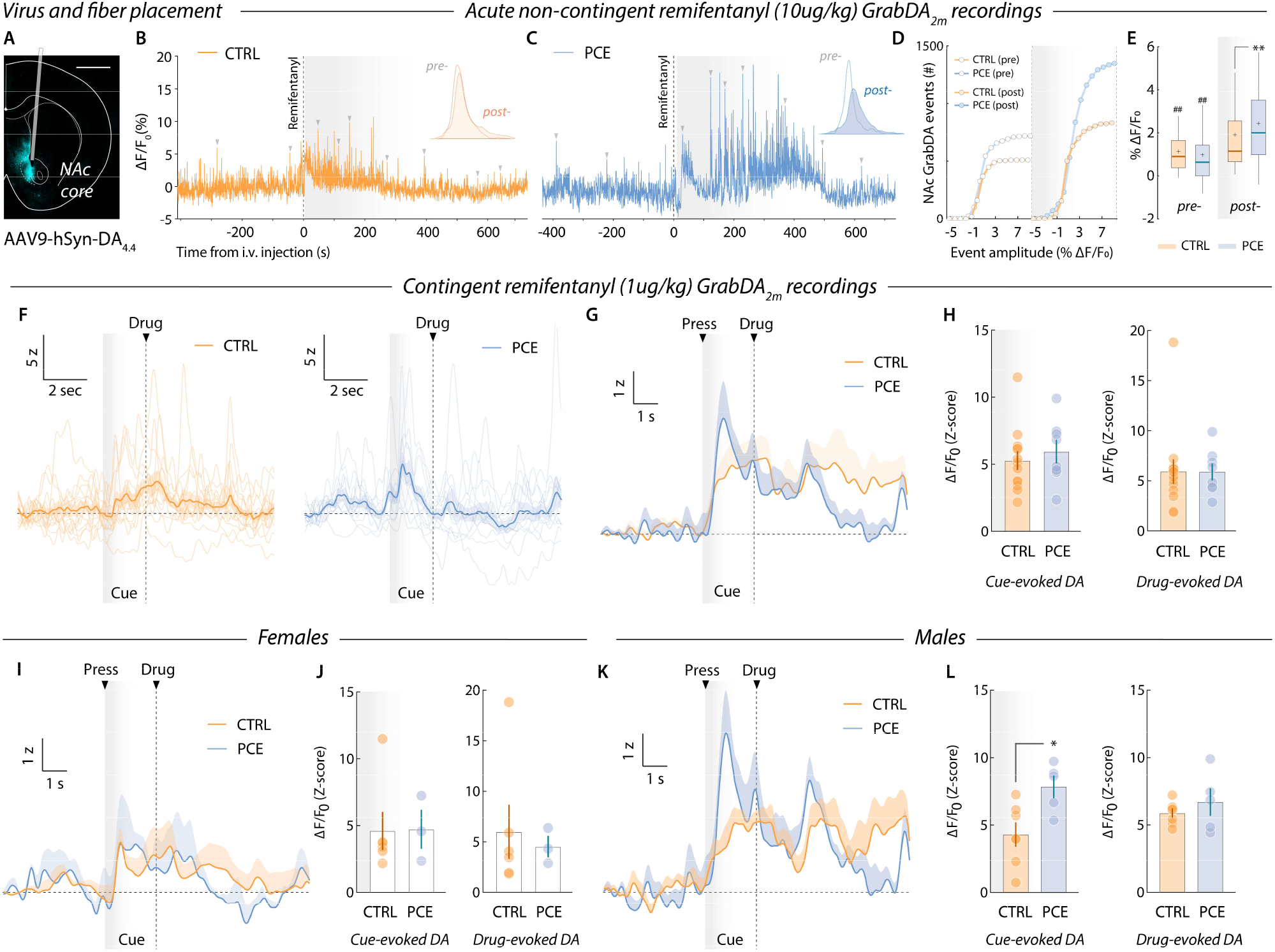
NAc dopamine release dynamics during remifentanyl injection and voluntary intake. **A**) Representative image showing viral expression localization and optic fiber placement in the NAc core. **B**,**C**) Spontaneous GrabDA_2m_ transients (indicated with grey triangles) before and after remifentanil i.v. injection by experimenter. Insets illustrate distributions of spontaneous event amplitudes (% ΔF/F_0_) pre-(white) and post-injection (colored). **D**) Cumulative amplitude distribution of NAc dopamine release events before and after remifentanyl administration. **E**) Pre-post injection comparison of NAc dopamine events from control and PCE rats (two-way RM ANOVA; ^*treatment*^*F*_1,3462_ = 4. 37, *p* = 0.030; ^*remifentanyl*^*F*_1,3462_ = 176.8, *p* < 0.001; ^*interaction*^*F*_1,3462_ = 85.5, *p* < 0.001 –followed by Bonferroni’s post-hoc: pre-post CTRL, ^##^*p* < 0.001; pre-post PCE, ^##^*p* < 0.001; Post-, CTRL vs. PCE, ^**^*p* < 0.001). *n*_CTRL_ = 5; *n*_PCE_ = 4. Center line represents the median; the cross illustrates the average; the bounds of the box depict the 25^th^ to 75^th^ percentile interval; and the whiskers represent minima and maxima. **F**) Representative trial-by-trial and average NAc dopamine transients in response to drug-paired cue presentation during PR. **G**) Group-averaged NAc dopamine transients. **H**) Cue-(0-2sec, relative to cue onset) (*t*_19_ = 0.63, *p* = 0.53) and drug-evoked (2-8sec, relative to cue onset) (*t*_19_ = 0.02, *p* = 0.97) dopamine amplitudes (AUC) in control and PCE rats (*n*_CTRL_ = 13; *n*_PCE_ = 8). **I**,**J**) Group-averaged NAc dopamine PSTHs from control and PCE female animals (cue-evoked amplitudes: *t*_7_ = 0.05, Bonferroni-Dunn-corrected *p* > 0.99) (drug-evoked amplitudes: *t*_7_ = 0.37, Bonferroni-Dunn-corrected *p* > 0.99). Females, *n*_CTRL_ = 6; *n*_PCE_ = 3. **K**,**L**) Group-averaged NAc dopamine PSTHs from control and PCE male rats (cue-evoked amplitudes: *t*_10_ = 2.89, Bonferroni-Dunn-corrected *p* = 0.031) (drug-evoked amplitudes: *t*_10_ = 0.91, Bonferroni-Dunn-corrected *p* = 0.76). Males, *n*_CTRL_ = 7; *n*_PCE_ = 5. For all bar and point plots, bars denote mean ± SEM.

### Apparatus

Rats were tested in operant chambers (12.0’’ L x 9.5’’ W x 8.25’’, Med Associates) inside sound-attenuating cabinets. Each chamber was equipped with two retractable levers (located 2 cm above the floor) as well as one LED stimulus light and a 2.5 kHz tone-generating speaker located above each lever. An external food magazine delivered food pellets to a dispenser centrally located between the two levers. Alternatively, remifentanyl infusions were delivered in a 20 ml injection over 2-s via a syringe mounted on a microinfusion pump (PHM-100A, Med-Associates) through a single-channel liquid swivel (375/25, Instech Laboratories) connected via tygon tubing to the chronic indwelling catheter. A house light and a white-noise speaker (80 dB, masking noise background) were located on the opposite wall. Throughout all operant paradigms, a multi-sensorial combination of conditioned stimuli (lever retraction, light stimuli, auditory cues) was used to avoid potential sensorial confounds imposed by PCE.

### Food self-administration

After 2-4 weeks of recovery and viral expression, or ∼PND120, rats were food-restricted to 85-90 % of their original bodyweight and trained to lever press for palatable food pellets (Bio-Serv #F0299) under a FR1 schedule of reinforcement. After nine 45-min daily sessions, rats progressed to a FR5 schedule, which continued for six days. Following FR5 training, a PR schedule of appetitive reinforcement was used to estimate the effort rats were willing to expend for a food reward. On each successive trial, the response requirement (lever presses) needed to obtain reward scaled near-logarithmically. The response ratio of the first sixteen trials was: 1, 2, 4, 6, 9, 12, 15, 20, 25, 32, 40, 50, 62, 77, 95, 118. The last response requirement attained, also known as break-point, was recorded and used to infer the inherent motivation for the reward. The PR session ended whenever no reward could be obtained within 10 min. During FR1, FR5 and PR training, animals had continued access to the active and inactive levers (except for the timeout period, 2-s) and each active lever press was rewarded with a single pellet. Responses on the inactive lever were recorded but had no programmed consequences.

### Exponential food demand task

Following PR testing, a subset of food-trained rats were tested on five 50-min demand task sessions. Each session was divided into five 10-min bins in which the lever pressing requirement for earning a food pellet followed a descending trend: 180, 90, 45, 15, 5. We used a descending unit price to prevent excessive food intake and satiation during the beginning of the session. We recorded the number of pellets in each bin (averaged across sessions) as the primary dependent measure during this experiment. Inactive lever presses had no consequences. We fitted the data to the exponential model of demand equation^45^:

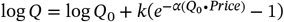

Where *Q*_*0*_ is maximal demand at zero cost, *k* is a shared scaling constant that reflects the range of the data, *α* determines the rate of decline in relative consumption and ‘*price*’ is the independent variable of cost (responses required per pellet). We calculated essential value (EV) and *P*_*max*_ for each animal using the following formulas^82^:

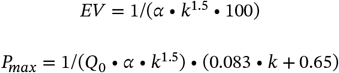

### Go/No-go task

Following behavioral economics testing, rats started go/no-go pretraining. In this phase (3 daily sessions), rats were only cued with go trials (cue light, 2.5-kHz tone), in which they were required to lever press within 5-sec of its presentation in order to obtain a pellet reinforcer. If the 5 sec elapsed with no response, the lever would retract, no reward would be presented, and a new trial would begin. Inter-trial interval (ITI) was on average 40-sec, and each session consisted of 64 trials. Rats were then trained to respond to both go and no-go trials (white noise, house light). 32 trials of each class were presented each session, and their order was pseudorandomized in blocks of 10-11 trials. Go and no-go trials were cued during 5-sec. After thirteen sessions, rats progressed to phase 2 of go/no-go responding, in which all parameters were kept the same but no-go trials were extended to a 10-sec duration.

### Intravenous remifentanyl self-administration

After 2-4 weeks of recovery and viral expression, or ∼PND115, rats were subjected to jugular vein catheterization surgery. Surgical implantation of the catheter was performed following anesthesia using isoflurane in O_2_ (4% induction and 2% maintenance). Indwelling i.v. silastic catheters (1.6 mm outer diameter) (Instech #C30PU-RJV1611) were implanted 3.8 cm into the right jugular vein and anchored with suture. The remaining tubing ran subcutaneously to the 22 ga vascular access button (Instech #VABR1B/22), which exited at the midscapular region and remained protected with a magnetic aluminum cap (Instech #VABRC). All incisions were sutured and coated with antibiotic ointment (Bactroban, GlaxoSmithKline). After surgery, animals were allowed to recover for 7-5 days prior to initiation of self-administration sessions. To maintain patency, catheters were flushed daily with heparinized saline (30 USP units/ml). After surgery recovery, rats were trained (FR1) to lever press for remifentanyl infusions (0.9 μg/kg in 20 μl injection over 2-sec) through a single-channel liquid swivel (Instech Laboratories #375/22PS) connected via tygon tubing to the chronic indwelling catheter. After seven sessions, response requirements increased (FR5) and testing continued for five sessions more. Each infusion was followed by a 10-sec time-out period in which a nosepoke on the active hole had no consequences but was recorded. Then, rats were tested in a PR schedule for 3 daily sessions (as previously explained). After each experimental phase transition, rats showing marked signs of body weight loss or distress were removed from the study.

*Extinction and cue-induced reinstatement*: following remifentanyl intake phases, rats underwent operant extinction of drug-seeking behavior. Extinction sessions (2-h long, once daily) were conducted for a minimum of seven days and active lever presses produced neither drug delivery nor cue presentation. When animals showed less than 30 lever presses for two consecutive extinction sessions, they were re-introduced to the drug-associated compound cue. The cue-induced reinstatement sessions (FR1) lasted 2-h and lever presses resulted in cue presentation and microinfusion pump activation, but not remifentanyl infusion.

### Exponential remifentanyl demand task

Following PR responding, a subset of remifentanyl-trained rats were tested on an exponential remifentanyl demand task. All training and modeling parameters were the same as in the food demand task, with the exception of the pressing requirement progression, which was adjusted to: 6, 10, 16, 25, 40, 63, 100, 158^83^ and resulted in 80-min sessions.

### Fiber photometry

Four weeks before their respective operant training procedures started, food- and remifentanyl-seeking rats were subjected to chronic optical fiber implantation surgery. Rats were anaesthetized with isoflurane in O_2_ and then placed in the stereotaxic apparatus. The viral vector AAV9-hSyn-DA-sensor 4.4 (GrabDA_2m_) (1 μl, 10^13^ gc/ml) was unilaterally injected in the NAc using the following coordinates: AP +1.3 mm, ML ±1.4 mm and DV -7.2 mm, relative to bregma (skull surface). Immediately following virus infusion, an optical fiber (diameter, 400 μm; NA, 0.5; Thorlabs) embedded within a ceramic ferrule was implanted with its tip targeting 0.1 mm above the above-mentioned NAc DV coordinates. Fiber photometry of GrabDA_2m_ signals was conducted in all PR sessions. Two fiber-coupled LEDs producing 470 nm and 405 nm lasers (Lx465, Lx405; Tucker-Davis Technologies, TDT) were used as the excitation source. The LED beams were reflected and coupled to a fluorescence minicube (FMC4, Doric Lenses). A 2 m-long optical fiber (400 μm, Doric Lenses) was used to transmit light between the fluorescence minicube and the implanted fiber. A RZ10x real-time processor (Tucker-Davis Technologies, TDT) was used to convert the current signal to voltage signal, which was processed through a low-pass filter (6Hz, sixth order Butterworth filter) built-into the Synapse software (TDT). GrabDA_2m_ signals were processed using the MATLAB script developed by Barker et al.^84^. Noise-related changes in fluorescence across the whole experimental session were removed by scaling the isosbestic control signal (405 nm) and regressing it onto the dopamine-sensitive signal (470 nm). Z-scored PSTHs were obtained after each PR ratio completion (baseline -4 to -2 sec, relative to cue onset). Alternatively, spontaneous GrabDA_2m_ transients before and after the remifentanyl (10 μg/kg, i.v.) injection test were identified using the ‘*findpeaks*’ (min peak prominence = 1) MATLAB function.

### *In vivo* electrophysiology

Two weeks before food or remifentanyl operant training, rats underwent surgical implantation of an 8 or 16 microwire array. Electrodes were made of Teflon-insulated stainless steel (0.25 mm interelectrode space, 0.5 mm inter-row space; Micro Probe). Electrodes were fixed to the skull with acrylic cement and stainless-steel bone screws. A stainless-steel wire accompanying each array served as a ground electrode and was inserted into the midbrain/cerebellum. The Multi-channel Acquisition Processor (Plexon Inc.) was used for *in vivo* electrophysiology recordings. Headstages for either 8 or 16 electrodes with 1X gain and 36-gauge wire cables were used. A second level of amplification occurred at a 16-channel differential preamplifier with a fixed gain of 100X.

Further amplification was performed through the online sorter software (gain 1000-32000), filtering with a 500 Hz high pass cut off and a bandpass filter between 0.7 Hz-300 Hz. Thresholds for each channel were chosen by the experimenter and typically 1 to 5 units were recorded per channel, excluding the reference channel. Voltage boxes were used to sort spikes within a channel. Activity was recorded for a three-minute baseline before the start of behavioral paradigms and terminated at the completion of the task. Offline spike sorting was performed in Offline Sorter (Plexon Inc) based on PCA and visual inspection of waveforms. After the spike-sorting procedure, the firing rate of each neuron was normalized to the average firing rate for that neuron during the entire session. Normalized firing rates were then standardized (z-scored) across units. After normalization, a Gaussian kernel sliding window of 8 bins (σ = 3) was applied (‘*gaussfilter*’ MATLAB function). Distinct NAc patterns of encoding during PR were uncovered using supervised *k*-means clustering. The features extracted for clustering were: (*1*) number of transients (peaks and troughs), (*2*) time to first transient event relative to cue onset, (*3*) mean activity during cue presentation and (*4*) mean activity after reward delivery. *k* values (number of clusters) were iteratively increased as long as new encoding patterns emerged. The final *k* value corresponded to the iteration prior to finding that the new patterns discovered resembled those already existing (*k* = 4 for the food PR dataset, *k* = 3 for the remifentanyl PR dataset). Neurons that did not cluster into any pattern were considered as ‘non-encoding’. Mean normalized firing rates (AUCz) were obtained using the area under the curve (‘*trapz*’ MATLAB function) during the corresponding analysis window (cue: 0-2 sec, reward: 2-7 sec; relative to cue onset, except otherwise stated).

### Tensor Component Analysis (TCA)

To find a compact and interpretable description of the multi-trial neural activity obtained from the PR task, we perform a dimensionality reduction technique called canonical polyadic (CP) tensor decomposition^85^. CP is a generalization of PCA to higher-order data arrays (tensors). In this case, we fitted the dimensions *U × T × P*, where *U* are the individual recorded units, *T* are PR completed trials and *P* is the firing rate PSTH around each cue presentation. Based on Williams et al. (2018)^27^ study and MATLAB package (https://github.com/ahwillia/tensortools) we used the *alternating least-squares* (ALS) algorithm to iteratively optimize dimensionality reduction solutions until converging to minimal reconstruction errors. ALS optimizes one of the factors (e.g., the unit factor W_u_) while fixing the other two (the PSTH and trial factors A_p_ and B_t_, respectively), yielding the following update rule:

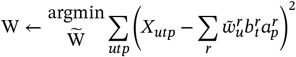

This procedure is then cyclically applied by optimizing one of the other factors and fixing the remaining two. Fitting TCA to data is a nonconvex problem^27^. Thus, the model was fitted multiple times, each time starting from a different random initial parameter set, and the number of components, here referred to as ‘cell assemblies’, was gradually adjusted until the smallest number of recovered components (*r = 4*) that converged across multiple runs was found. Importantly, these low-dimensional four cell assemblies largely resembled those recovered by traditional *k*-means clustering. To ensure all arrays (dimensions) were of the same length, only data from the first 8 PR trials (lowest break-point registered) was included. Functional relationships between uncovered cell assemblies with response vigor during PR was examined using a machine learning prediction algorithm. A gradient boosted trees model was trained on each neuron unit factors W_u_ (80% data hold for the training set) to predict latency to 8^th^ PR reward (TensorFlow, Google). The validity of this prediction was determined by correlating ground truth values with predicted latencies (Pearson’s *r*).

### Histology

Rats underwent isoflurane anesthesia (5%) and transcardial perfusion with a 4% paraformaldehyde (PFA) solution in a 0.1M sodium phosphate buffer (PB) at pH7.4. Following perfusion, brains were post-fixed at 4 °C in PFA overnight. Brain sections, (40 µm thick) were obtained using a vibratome (Leica). Coronal sections were immersed in a PB solution containing 3% normal donkey serum (Jackson 017-000-121) for a duration of 30 min. Immunodetection of GrabDA_2m_ was achieved following 2-hour incubation with an anti-GFP antibody (ThermoFisher #A-11122) (1:200) and labeled with goat anti-rabbit secondary antibody Alexa Fluor Plus 555 (ThermoFisher #A32732) (1:2000, 45 min).

## Statistical analyses

Parametric behavioral, dopaminergic, and firing rate (AUCz) measures were analyzed using a one- or two-way repeated measures (RM) ANOVA, or unpaired two-tailed Student’s *t* tests and Bonferroni post hoc tests were used to correct for multiple comparisons when appropriate. Every time samples were split into different groups, such as sexes, following Student’s *t* tests were corrected for multiple comparisons using the Bonferroni-Dunn method. Whenever violations of homoscedasticity were detected, a Welch correction was applied (Welch’s *t* test). When analyzing non-parametric variables (proportion of subjects), the *Chi-squared* tests were employed. Significance was set at *p* < 0.05 and all tests were two-tailed. To determine biases in the assignment of neurons corresponding to the identified clusters (Figure **3** and **6**), a Bayesian logistic regression model using a compound confluent hypergeometric prior distribution (*α* = 0.5, *β* = 2, *s* = 0) was constructed. Statistical analyses were performed in MATLAB 2023b, GraphPad Prism 10.1 and JASP 0.18.

**Figure 6.**
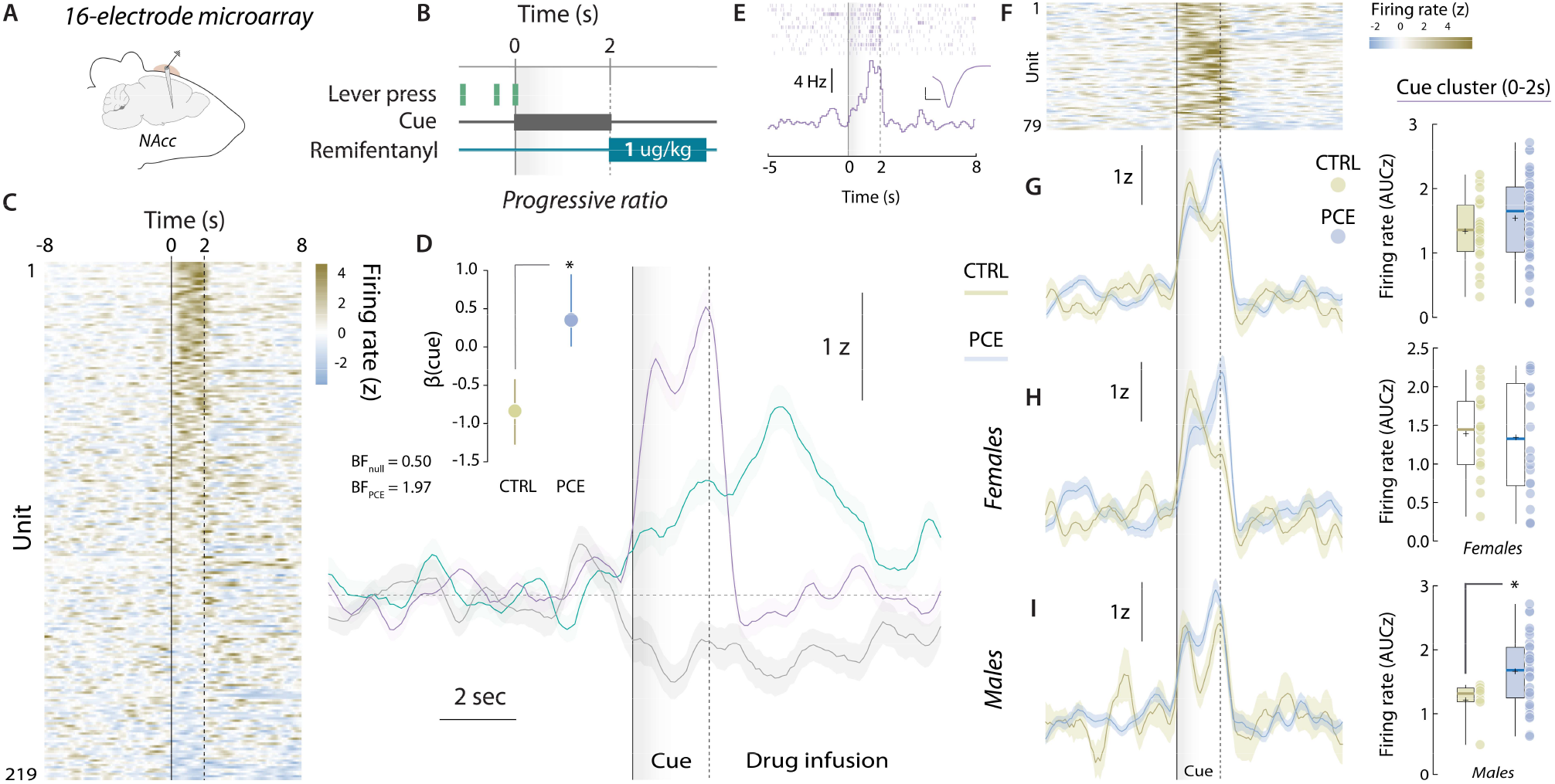
NAc encoding of drug-paired cues during voluntary remifentanyl pursuit. **A**,**B**) Schematic representation of chronic multi-electrode microarray NAc implantation and PR operant task structure. **C**) Trial-averaged, cue-aligned PSTHs of all NAc units recorded during PR responding. **D**) Trial-averaged firing rates from the cue-responding (*n*_CTRL_ = 23; *n*_PCE_ = 56), reward-responding (*n*_CTRL_ = 23; *n*_PCE_ = 39) and lever press-inhibited clusters units (*n*_CTRL_ = 31; *n*_PCE_ = 23). (Inset) Posterior coefficients (β) with credible intervals (vertical lines) of Bayesian logistic regression models evaluating the influence of PCE on the likelihood to find a unit on the cue-responding cluster. Bayes factors (BF) for null and PCE-influenced hypotheses are shown as insets. ‘*’ Represents ‘anecdotic’ evidence in favor of the alternative (PCE-influenced) hypothesis. **E**) Raster (top) and PSTH (bottom) plot of a representative neuron pertaining to the cue-responding cluster. Inset panel shows the corresponding waveform (scale bar: 50 μV, 20 ms). **F**) Trial-averaged heatmap depicting all neurons’ activity from the cue-responding cluster. **G**) (Left) Trial-averaged PSTH of the cue-responding cluster in control and PCE rats. (Right) Average firing rates (AUC) during cue presentation (grey shaded area, 0-2sec) from cue-responding units (*t*_77_ = 1.36, *p* = 0.17). **H**,**I**) (Left) Cue-responding cluster neurons average firing rates from female and male animals. (Right) Average firing rates (AUC) during cue presentation from cue-responding female (*t*_36_ = 0.21, Bonferroni-Dunn-corrected *p* > 0.99) and male neurons (\**t*_39_ = 3.01, Bonferroni-Dunn-corrected *p* = 0.018). *n*_CTRL_ = 23 (16F, 7M); *n*_PCE_ = 56 (22F, 34M). Center line represents the median; the cross illustrates the average; the bounds of the box depict the 25^th^ to 75^th^ percentile interval; and the whiskers represent minima and maxima.

**Figure 7.**
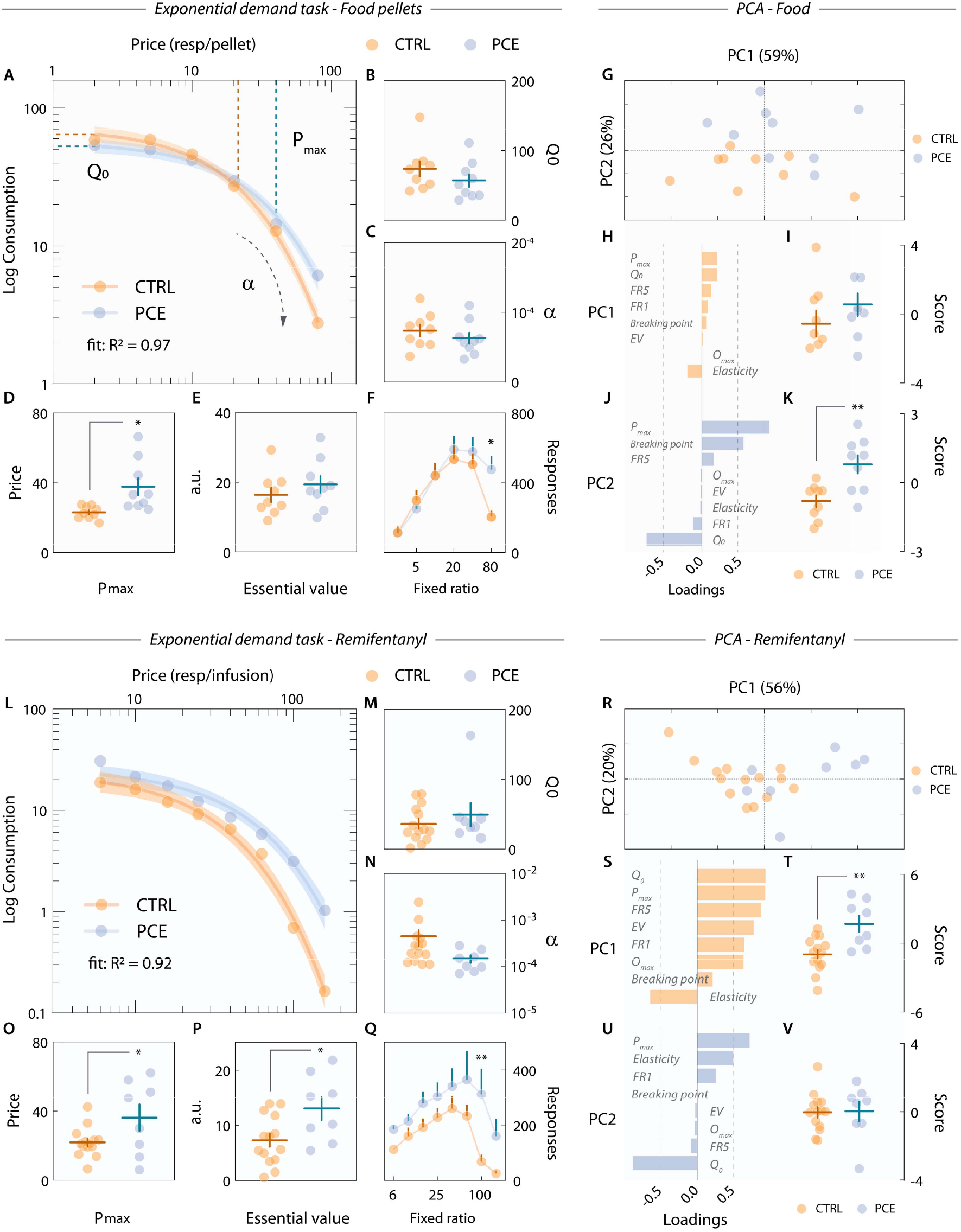
Behavioral economics modelling of food and opioid (remifentanyl) demand. **A**) Exponential demand curve for palatable pellet rewards. Colored shaded areas represent 95% CI. **B**) *Q*_*0*_, the theoretical demand at a unit price of 0 lever presses, in control and PCE rats (*t*_16_ = 1.21, *p* = 0.24). **C**) PCE does not influence price sensitivity (α) (*t*_16_ = 0.94, *p* = 0.36). **D**) PCE rats are willing to pay higher prices (*P*_*max*_) for the same valued reward than control subjects (Welch’s \**t*_9.0_ = 2.92, *p* = 0.017). **E**) Unchanged reinforcing efficiency of food pellets (essential value) in control and PCE rats (*t*_16_ = 0.97, *p* = 0.34). **F**) Session-averaged lever press responses at each price point (two-way RM ANOVA; ^*treatment*^*F*_1,16_ = 0. 82, *p* = 0.37; ^*price*^*F*_5,80_ = 28.5, *p* < 0.001; ^*interaction*^*F*_5,80_ = 3.13, *p* = 0.012 –followed by Bonferroni’s post-hoc: CTRL vs. PCE, \**p* = 0.013). **G**) Individual scores on the two principal components identified by PCA. PC1: ‘inelastic lever pressing’, PC2: ‘anhedonic motivational drive’. Percentages indicate the proportion of total behavioral variance explained by each factor. **H**) Variable loadings onto PC1. **I**) Control and PCE rat ‘inelastic lever pressing’ scores (*t*_16_ = 0.97, *p* = 0.34). **J**) Variable loadings onto PC2. **K**) Control and PCE rat ‘anhedonic motivational drive’ scores (\*\**t*_16_ = 3.46, *p* = 0.003). For all food pellet panels, *n*_CTRL_ = 9; *n*_PCE_ = 9. **L**) Exponential demand curve for remifentanyl. Colored shaded areas represent 95% CI. **M-P**) *Q*_*0*_ (*t*_20_ = 0.85, *p* = 0.40), α (Welch’s *t*_13.6_ = 1.78, *p* = 0.097), *P*_*max*_ (\**t*_20_ = 2.23, *p* = 0.037) and essential value (\**t*_20_ = 2.57, *p* = 0.018) derived from control and PCE animals’ remifentanyl demand curves. **Q**) Session-averaged lever presses for remifentanyl at each price point (two-way RM ANOVA; ^*treatment*^*F*_1,20_ = 4.64, *p* = 0.043; ^*price*^*F*_7,140_ = 13.7, *p* < 0.001; ^*interaction*^*F*_7,140_ = 2.37, *p* = 0.025 –followed by Bonferroni’s post-hoc: CTRL vs. PCE, \*\**p* = 0.001). **R**) Individual scores on the two principal components identified by PCA in the remifentanyl dataset. **S**) Variable loadings onto PC1. **T**) Control and PCE rat ‘inelastic lever pressing’ for remifentanyl scores (\*\**t*_20_ = 3.53, *p* = 0.002). **U**) Variable loadings onto PC2. **V**) Control and PCE rat ‘anhedonic motivational drive’ for remifentanyl scores (*t*_20_ = 0.09, *p* = 0.92). For all drug-related panels, *n*_CTRL_ = 14; *n*_PCE_ = 8. For all bar and point plots, lines denote mean ± SEM.

**Figure 8.**
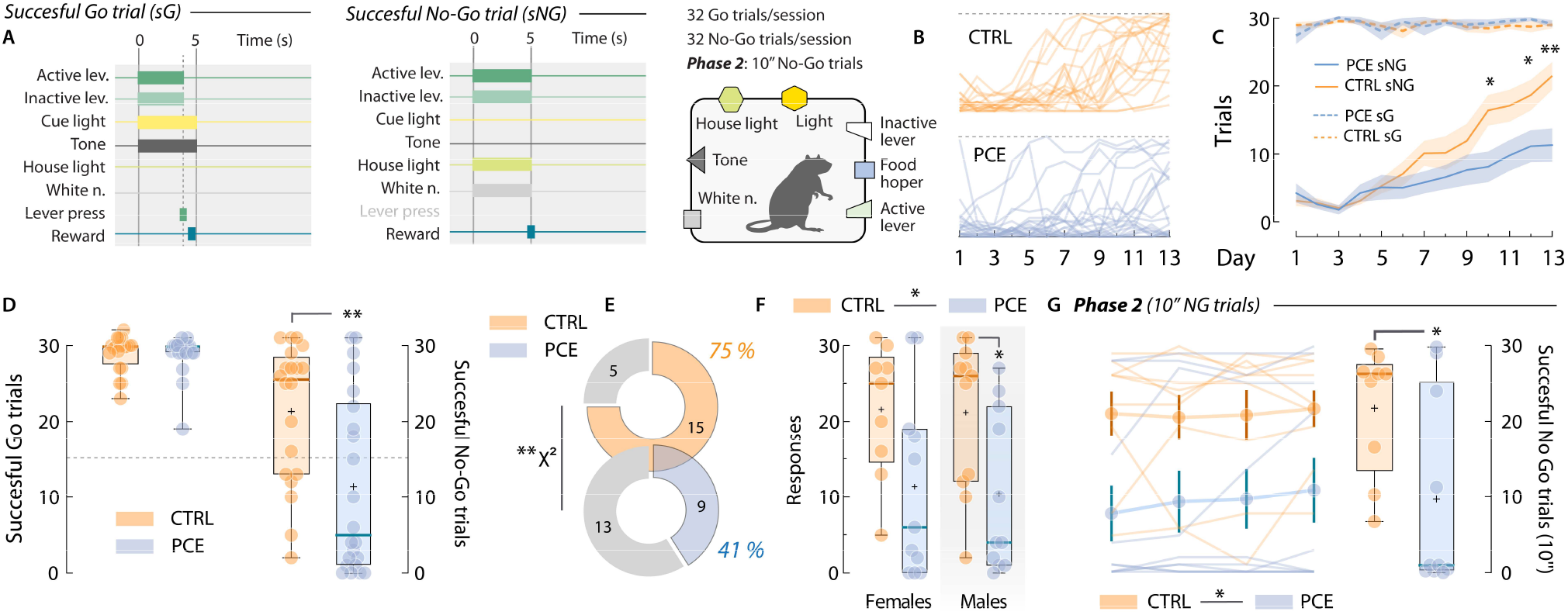
PCE increases impulsive behavior in a go/no-go task. **A**) Schematic illustrating the compound cues and manipulanda available in the operant chamber during inter-trial intervals, as well as the consequences (pellet delivery) of successfully executed go and no-go trials. **B**) Individual daily progression of successful no-go responding in control and PCE rats. **C**) Group-averaged number of successfully executed go (sG, dashed lines) and no-go (sNG, solid lines) trials (two-way RM ANOVA; ^*treatment*^*F*_1,40_ = 3.36, *p* = 0.07; ^*day*^*F*_12,480_ = 29.6, *p* < 0.001; ^*interaction*^*F*_12,480_ = 4.65, *p* < 0.001 –followed by Bonferroni’s post-hoc: CTRL vs. PCE; day 10 \**p* = 0.016, day 12 \**p* = 0.016, day 13 \*\**p* = 0.001). **D**) Successfully executed go (*t*_40_ = 0.17, *p* = 0.86) and no-go trials (\*\**t*_40_ = 3.07, *p* = 0.003) after training (day 13). Dashed line marks 50 % performance. **E**) Percentage of control and PCE rats obtaining more than 50 % of no-go trial rewards (χ^2^ = 61.6, \*\**p* < 0.001). **F**) Successfully executed no-go trials by sex (females; *t*_18_ = 2.11, Bonferroni-Dunn-corrected *p* = 0.097) (males; *t*_20_ = 2.47, Bonferroni-Dunn-corrected \**p* = 0.045). *n*_CTRL_ = 20 (9F, 11M); *n*_PCE_ = 22 (11F, 11M). **G**) Daily (left) and average (right) successfully executed 10” no-go trials in phase 2 (\**t*_17_ = 2.38, *p* = 0.029) (*n*_CTRL_ = 9; *n*_PCE_ = 10). For all box plots, center line represents the median; the cross illustrates the average; the bounds of the box depict the 25^th^ to 75^th^ percentile interval; and the whiskers represent minima and maxima.

## Supplementary Figures

**Supplementary Figure 1.**
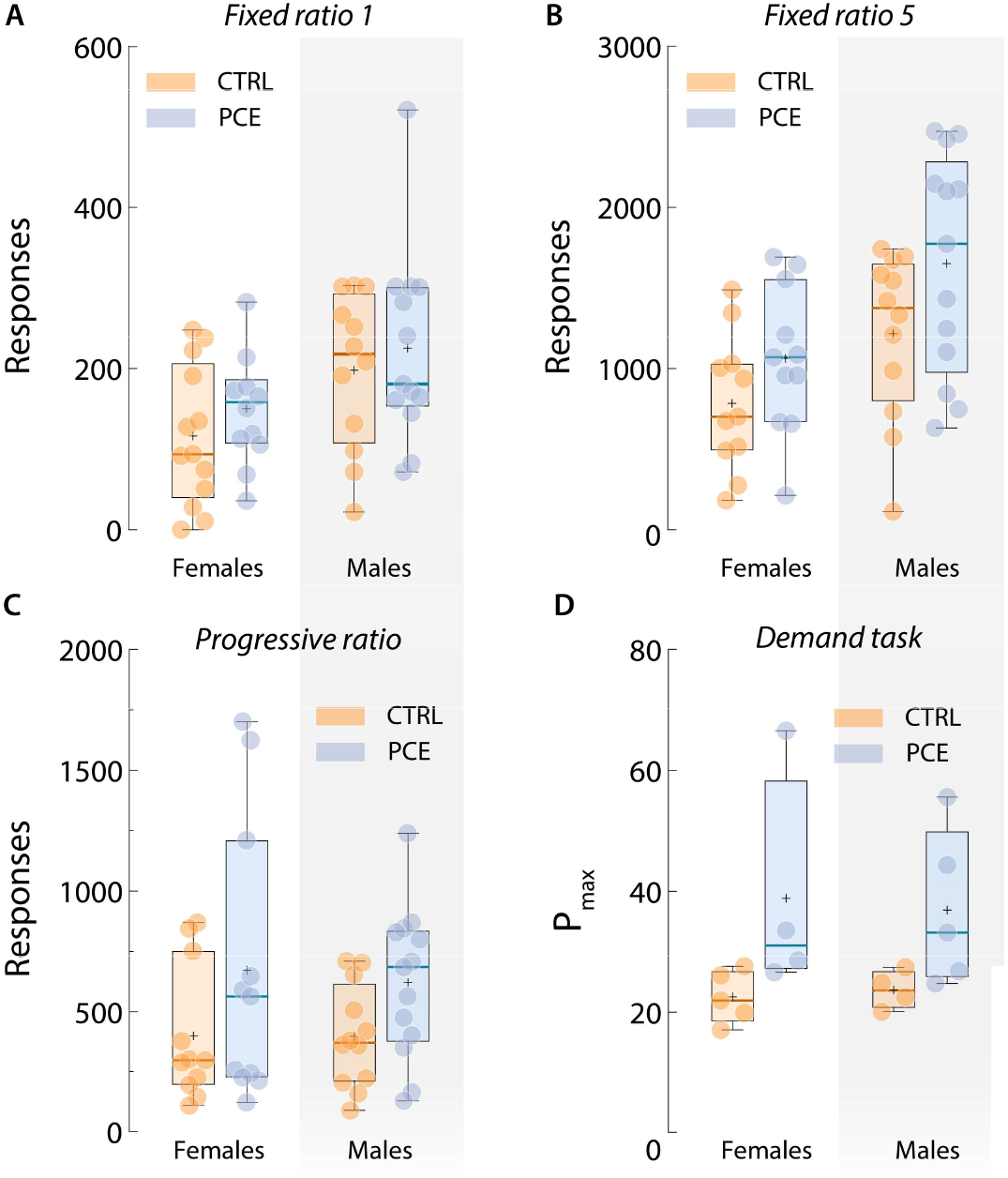
Lack of sex-dependent effects of PCE on motivation for food rewards. **A)** Total lever presses under a FR1 schedule of reinforcement (females; t22 = 0.99, Bonferroni-Dunn-corrected p = 0.66) (males; t23 = 0.54, Bonferroni-Dunn-corrected p > 0.99). nCTRL = 25 (13F, 12M); nPCE = 24 (11F, 13M). B) Total lever presses on FR5 (females; t20 = 1.49, Bonferroni-Dunn-corrected p = 0.30) (males; t23 = 1.78, Bonferroni-Dunn-corrected p = 0.17). nCTRL = 23 (11F, 12M); nPCE = 24 (11F, 13M). C) Total responses on PR testing (females; t20 = 1.40, Bonferroni-Dunn-corrected p = 0.17) (males; t22 = 2.07, Bonferroni-Dunn-corrected p = 0.050). nCTRL = 23 (11F, 12M); nPCE = 24 (11F, 13M). D) Pmax values derived from the food demand task (females; t7 = 1.91, Bonferroni-Dunn-corrected p = 0.19) (males; t7 = 1.98, Bonferroni-Dunn-corrected p = 0.17). nCTRL = 9 (5F, 4M); nPCE = 9 (4F, 5M). Center line represents the median; the cross illustrates the average; the bounds of the box depict the 25th to 75th percentile interval; and the whiskers represent minima and maxima.

**Supplementary Figure 2.**
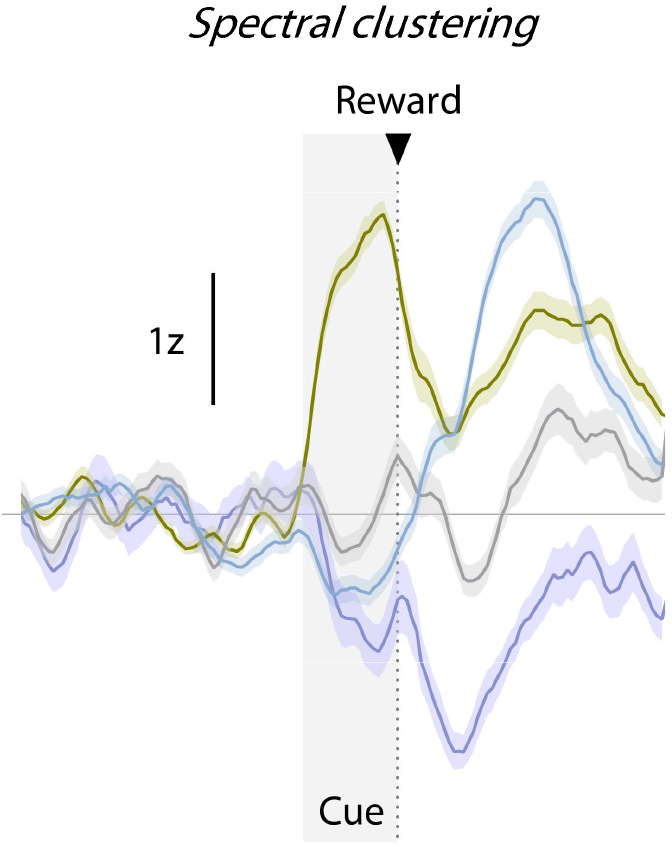
NAc patterns of encoding identified by spectral clustering. Trial-averaged, cue-Trialaveraged, cue-aligned PSTH of the four neuronal clusters uncovered by spectral decomposition. Lines represent average values and shaded areas ± SEM values.

**Supplementary Figure 3.**
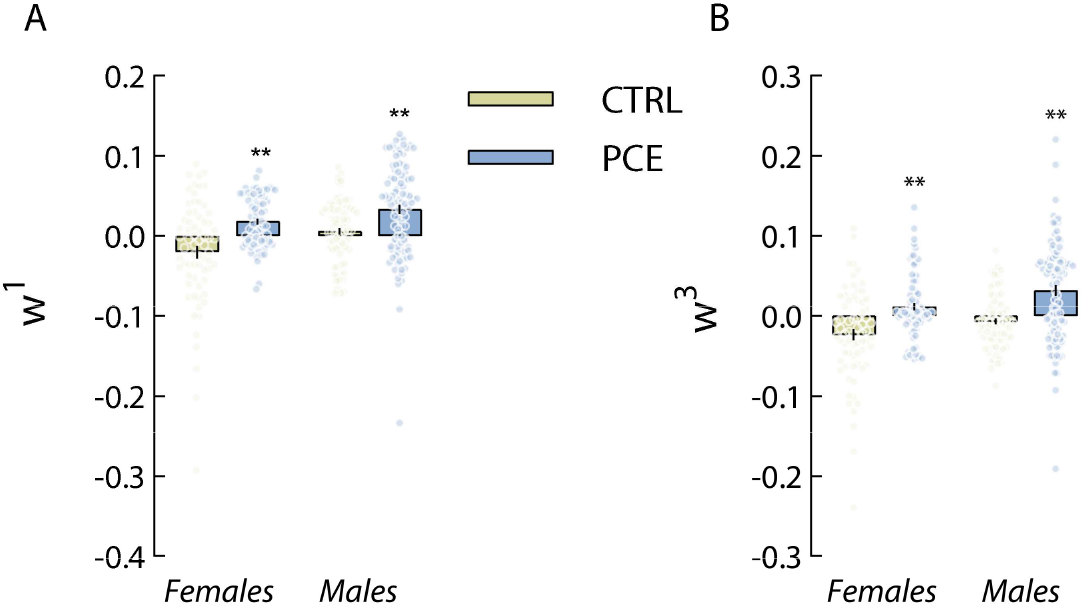
PCE effects on cue-encoding tensorial assemblies are not sex-dependent. **A)** Individual unit factor loadings (*w*^*1*^) of the cue-activated tensorial cell assembly (*a*^*1*^_*p*_*b*^*1*^_*t*_*w*^*1*^_*b*_) identified by TCA (females; *t*_143_ = 4.43, Bonferroni-Dunn-corrected \*\**p* < 0.001) (males; *t*_172_ = 3.50, Bonferroni-Dunn-corrected \*\**p* = 0.001). **B**) Individual unit factor loadings (*w*^*3*^) of the cue-inhibited tensorial cell assembly (*a*^*3*^_*p*_*b*^*3*^_*t*_*w*^*3*^_*b*_) (females; *t*_143_ = 4.21, Bonferroni-Dunn-corrected \*\**p* < 0.001) (males; *t*_172_ = 4.78, Bonferroni-Dunn-corrected \*\**p* < 0.001). *n*_CTRL_ = 143 (68F, 75M); *n*_PCE_ = 176 (77F, 99M). For all bar and point plots, bars denote mean ± SEM.

**Supplementary Figure 4.**
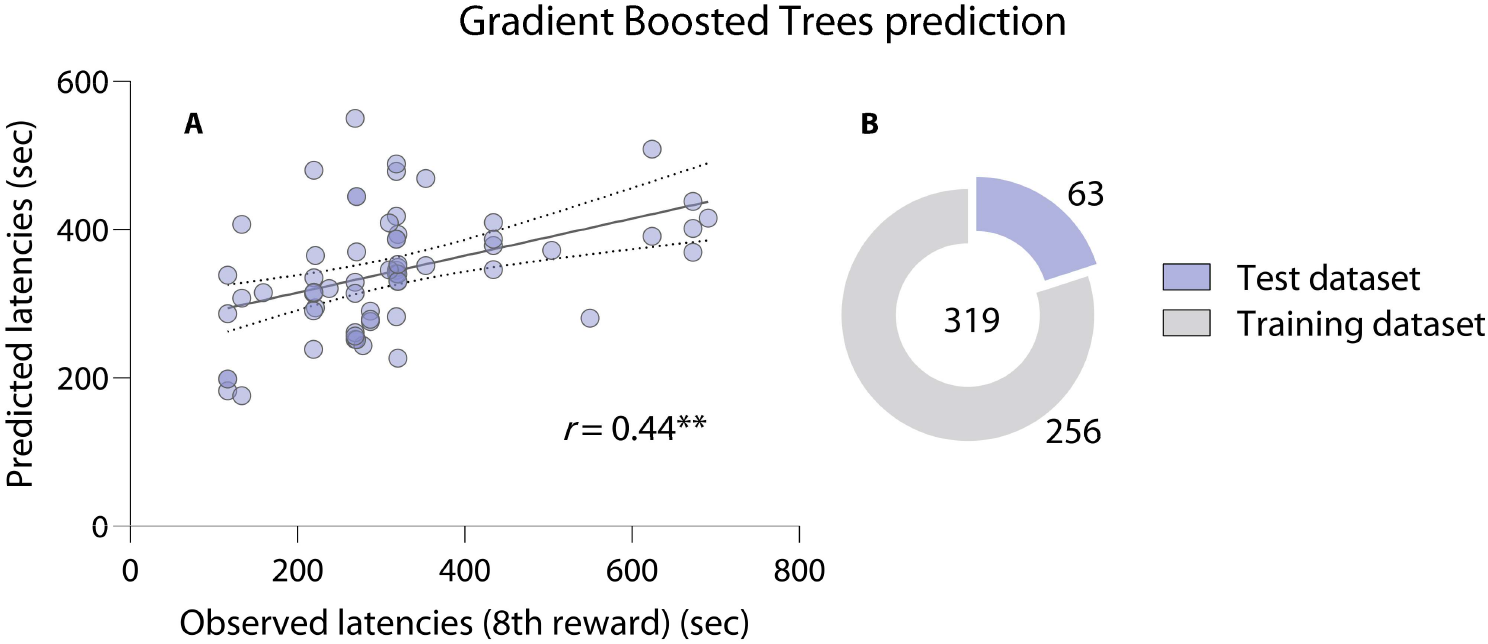
Tensorial unit loadings predict latency to reward hallmark during PR responding. **A)** Correlation between observed and predicted latency to complete 8 trials during PR (*ground truth* vs. *predicted value*; Pearson’s \*\**r* = 0.44, *p* = 0.003). **B**) A gradient boosted decision tree machine learning algorithm was trained on 80 % of the unit loadings and tested on the remaining 20 % cases.

**Supplementary Figure 5.**
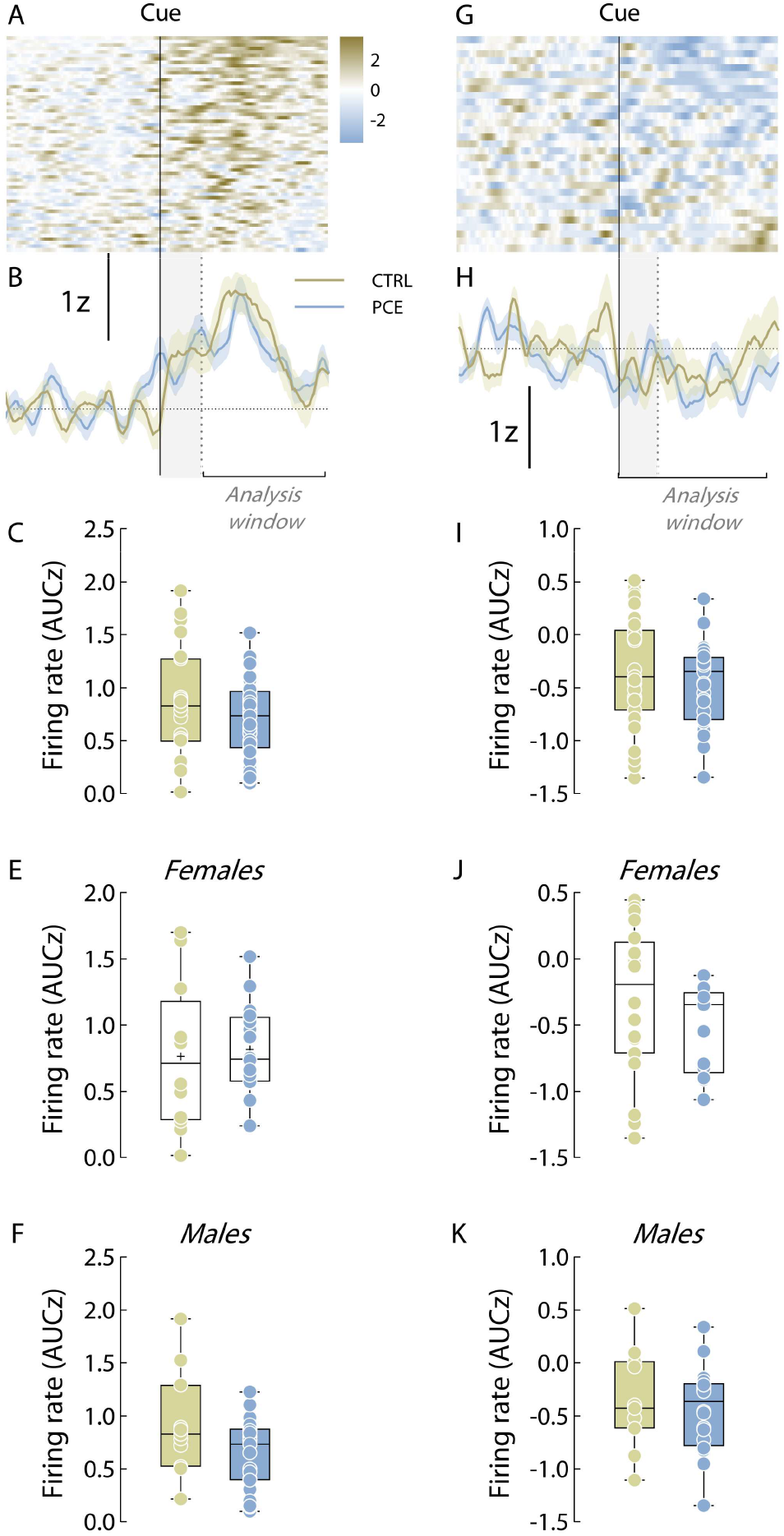
Reward-responding and lever press-inhibited NAc neuronal clusters identified during remifentanyl operant seeking. **A)** Trial-averaged heatmap depicting activity from all reward-responding units, centered around cue onset (vertical black line). B) Reward-responding neurons firing rate (Z-score). Shaded grey areas represent cue presentation and horizontal brackets indicate the time window used for parametric analyses. Lines represent average firing rates and colored shaded areas depict ± SEM. C-F) Trial-averaged firing rates (AUC) from the reward-activated cluster following remifentanyl infusion (Welch’s t33.4= 1.03, p = 0.31) (females; t26 = 0.31, Bonferroni-Dunn-corrected p > 0.99) (males; t32 = 1.94, Bonferroni-Dunn-corrected p = 0.12). nCTRL = 23 (12F, 11M); nPCE = 34 (11F, 23M). G-H) Heatmap and line plots depicting averaged encoding pattern of the lever press-inhibited cluster. I-K) Trial-averaged firing rates (AUC) from the lever press-inhibition cluster preceding cue presentation (t62= 1.27, p = 0.21) (females; t62 = 1.26, Bonferroni-Dunn-corrected p = 0.41) (males; t29 = 0.51, Bonferroni-Dunn-corrected p > 0.99). nCTRL = 42 (31F, 11M); nPCE = 53 (33F, 20M).

